# Taking identity-by-descent analysis into the wild: Estimating realized relatedness in free-ranging macaques

**DOI:** 10.1101/2024.01.09.574911

**Authors:** Annika Freudiger, Vladimir M. Jovanovic, Yilei Huang, Noah Snyder-Mackler, Donald F. Conrad, Brian Miller, Michael J. Montague, Hendrikje Westphal, Peter F. Stadler, Stefanie Bley, Julie E. Horvath, Lauren J. N. Brent, Michael L. Platt, Angelina Ruiz-Lambides, Jenny Tung, Katja Nowick, Harald Ringbauer, Anja Widdig

**Author notes:** AF, VMJ and YH contributed equally as first authors. KN, HR and AW contributed equally as senior authors.

## Abstract

Biological relatedness is a key consideration in studies of behavior, population structure, and trait evolution. Except for parent-offspring dyads, pedigrees capture relatedness imperfectly. The number and length of DNA segments that are identical-by-descent (IBD) yield the most precise estimates of relatedness. Here, we leverage novel methods for estimating locus-specific IBD from low coverage whole genome resequencing data to demonstrate the feasibility and value of resolving fine-scaled gradients of relatedness in free-living animals. Using primarily 4-6× coverage data from a rhesus macaque (*Macaca mulatta*) population with available long-term pedigree data, we show that we can call the number and length of IBD segments across the genome with high accuracy even at 0.5× coverage. The resulting estimates demonstrate substantial variation in genetic relatedness within kin classes, leading to overlapping distributions between kin classes. They identify cryptic genetic relatives that are not represented in the pedigree and reveal elevated recombination rates in females relative to males, which allows us to discriminate maternal and paternal kin using genotype data alone. Our findings represent a breakthrough in the ability to understand the predictors and consequences of genetic relatedness in natural populations, contributing to our understanding of a fundamental component of population structure in the wild.

## Introduction

Biological relatedness, defined as the sharing of alleles that are inherited from a common ancestor within a population, is a key concept in a wide range of disciplines, including behavioral ecology, quantitative genetics, and evolutionary genetics^1^. For example, biological relatedness is needed to estimate trait heritability^2,3^, calculate inbreeding for species management and conservation^4^, and test hypotheses for trait evolution^5^. However, most estimates of relatedness, especially in free-ranging populations, are based on the expected mean value for a given pedigree relationship^6^ and ignore substantial variation in “realized relatedness” (i.e., the true level of allele sharing between two individuals) within kin classes. This variation arises because of stochastic recombination during meiosis, which affects realized relatedness for all dyads in a pedigree other than parent-offspring dyads^7–14^. For example, although the expected proportion of allele sharing among full siblings is 0.5 (identical to parent-offspring dyads) the values actually observed in a set of 4,401 human full sibling dyads range from 0.37 to 0.62 (SD = 0.04)^10,15^. Deviations from pedigree expectations can be further exacerbated if the founders are related or if a pedigree contains gaps that obscure true genetic relationships^16–21^.

In principle, genetic markers can therefore be used as an alternative to pedigrees, providing relatedness estimates that directly reflect allele sharing between pairs of individuals. Indeed, many studies have used genetic variation at short tandem repeats (STRs, also often referred to as microsatellites) to estimate relatedness and/or categorize dyads into pedigree kin classes^22,23^. Yet, the precision of these estimates is often compromised by the use of relatively few STR markers (often <20), resulting in substantial statistical uncertainty and mistakes in kin category classification^14,24,25^. For example, van Horn et al.^25^ showed that a considerable fraction of known kin dyads were classified as nonkin using this method, while dyads with no known pedigree relationships were sometimes classified as kin. Such errors can lead to erroneous conclusions about the impact of kinship on social behavior.

However, following in the footsteps of human genetics and model systems research, studies of natural populations are increasingly transitioning away from small STR-based data sets to large-scale panels of single nucleotide polymorphism (SNP) markers collected via array technology or high-throughput sequencing^26–33^. Given a sufficient number of SNPs, these data can provide more accurate estimates of relatedness than pedigrees^34,35^. Perhaps even more excitingly, whole genome resequencing (WGS) data sets offer the possibility of inferring the precise stretches of DNA that are identical-by-descent (IBD, i.e., near-identical DNA, except for *de novo* mutations, that two individuals inherited from a recent common ancestor; Fig. 1A). The total fraction of the genome IBD between two individuals is, in principle, the most precise method to estimate kinship^36–38^, as it corresponds to the exact fraction of shared DNA (i.e., realized relatedness)^10,39–43^. Because distant relatives share fewer and shorter IBD segments than close relatives (as more recombination separates them from their common ancestors)^44–47^, the number and size distribution of IBD segments also provides additional information about pedigree relationships beyond that contained in mean IBD alone. This information can be very fine-grained: for example, in cases where recombination rates differ in female versus male germline cells, the number of shared segments is also expected to differ between maternal and paternal kin of the same degree^48,49^. Theoretically, this difference provides the basis for differentiating whether two individuals are connected via female versus male shared ancestors (e.g., are maternal half siblings versus paternal half siblings)—an important distinction in studies of behavior, given frequent differences in kin preferences and kin recognition for maternal versus paternal kin (reviewed in ^50^).

**Figure 1:**
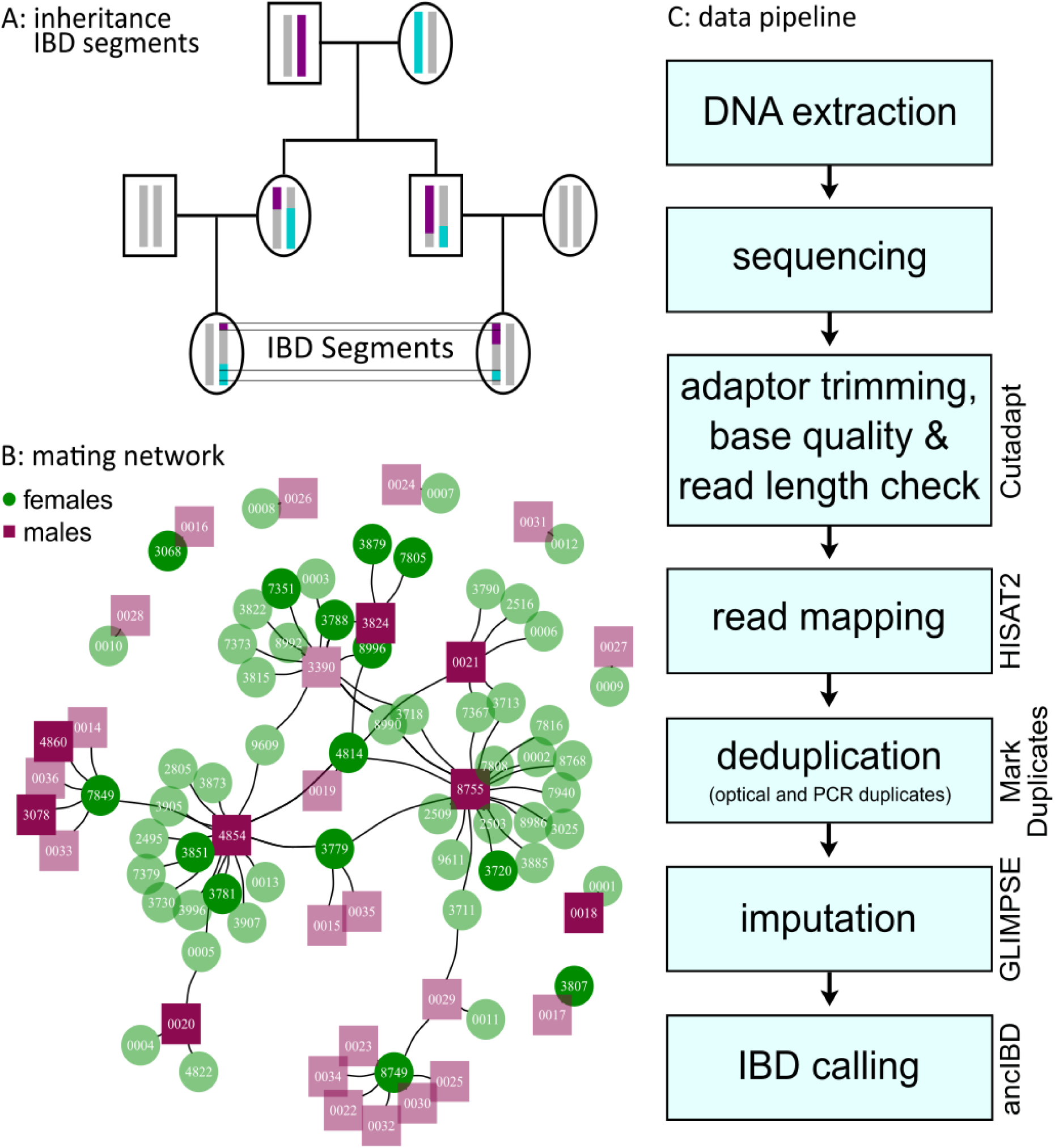
IBD inheritance, mating network and data pipeline. (A) Example of two IBD segments in full cousins, where blue segments are inherited from a common grandmother and purple segments are inherited from a common grandfather. (B) Mating network of the parents producing our study subjects. Females are shown as green circles, while males are shown as purple squares. Lines connect individuals that successfully reproduced together and thereby produced offspring included in this study. This illustrates that only a few very successful males have a large share of reproduction, while the differences in the number of reproductions are less pronounced in females. This mating network leads to a complex kinship structure, which is mainly based on half-siblings. Darker-colored individuals are parents that are included in the final data set. Due to space constraints, only the last four digits of the IDs are shown. (C) Data pipeline used for low coverage data. The names of the programs used are on the right of the box of each respective step.

Several methods are available to computationally infer IBD segments from high-quality genotype data^36–38,51^. In humans, these approaches have been used for a wide range of applications, including the detection of previously unknown biological relatives, reconstruction of historical patterns of population movement and admixture^52–54^, and the investigation of founder events that influence rates of genetic disease^55,56^. For example, IBD-based methods enable precise detection of relatedness structure, migration, and mating patterns even in previously unstudied populations^57,58^. However, IBD-based methods have rarely been applied outside of humans (but see for dairy cows^59^, maize^60^, captive rhesus macaques^47^, *Plasmodium*^61^). In natural populations, studies to date have typically either summarized IBD genome-wide or focused only on IBD within individuals (i.e., to identify runs of homozygosity indicative of inbreeding)^34,62–65^. While these initial successful applications are encouraging, they underscore the necessity for systematic testing on a population possessing a known pedigree to solidify this method as a reliable approach for estimating realized relatedness in nonhuman animals. They also relied on highly accurate genotype calls from moderate to high-coverage sequencing or other genome-wide genotyping platforms^47,62,63^. These approaches remain infeasible for many natural populations, where recent efforts to generate comprehensive genotyping data sets have focused instead on low-coverage resequencing^32,33^. However, advances in methods for genotype imputation suggest that locus-specific, population-scale IBD analysis may be possible even with these more limited data^66^. If so, highly accurate estimates of realized relatedness may be achievable in many more settings, including in long-term field studies where kin bias, inbreeding avoidance, and social evolution can be studied in nature.

Here, we pioneer such an approach to study realized relatedness in an intensively studied, free-ranging population of rhesus macaques (*Macaca mulatta*). To do so, we developed and tested a pipeline for low-coverage WGS data analysis and IBD calling by extending a method originally designed for ancient DNA studies in humans (*ancIBD*^67^). By combining our new approach with known pedigree data, state-of-the-art imputation methods, simulation, and downsampled high-coverage data, we demonstrate that precise and robust IBD calling is possible using WGS data at depths as low as ∼0.25 - 0.5× coverage. Further, we illustrate how IBD sharing can accurately measure biological gradients in pairwise relatedness, providing substantial information beyond the mean expected values obtained from pedigrees. Our findings point to several new insights obtainable from genome-wide IBD analysis, including the identification of cryptic relatives and empirical estimation of background levels of allele sharing even among putatively unrelated individuals. We also show that the number and length distribution of IBD segments are indeed sensitive to whether half sibling dyads share maternal versus paternal ancestry, despite no differences in the mean proportion of IBD genome-wide. Together, our results provide a broadly applicable roadmap for improving our understanding of genetic relatedness and its implications, including its role in social behavior, conservation, and evolutionary dynamics.

## Methods

### Study species and study population

The study population resides on Cayo Santiago, a 15.2 ha island off the southeast coast of Puerto Rico, USA, managed by the Caribbean Primate Research Center (CPRC). All monkeys living on the island descend from 409 founder animals that were captured at several different locations in the Lucknow area in northeastern India and brought to the island in 1938^68^. The distribution of annual reproduction among males on Cayo Santiago tends to be skewed, with only a limited number of males reproducing^69–71^. Therefore, half siblings are common while full siblings are rare^69,72^. Even though no new individuals have been introduced since 1938, available multi-generational pedigree data indicate a low incidence of inbreeding in this population^73^. To maintain the population size and genetic diversity at a sustainable level, the population is managed through regular removal of targeted individuals or groups^74^. The population is provisioned daily with monkey chow, but around one-fifth of their diet is based on natural vegetation^75^.

### Demographic and genetic data

Trained CPRC staff have collected census data since 1956, typically five days per week, including animal ID, date of birth and death, sex, maternal ID, and number of maternal kin^73^. Since 1992, biological samples have been collected from the vast majority of the population for paternity analyses using up to 42 STR markers^73,76^. The use of samples for this study was approved by the Institutional Animal Care and Use Committee (IACUC) of the University of Puerto Rico, Medical Science campus (protocol 4060105 to AW) and the transfer of samples followed the regulations of the Convention on International Trade in Endangered Species of Wild Fauna and Flora (CITES; US export: 22US26740E/9 and EU import DE-E-04076/22). Paternity is determined by a combination of genetic likelihood and exclusion analyses, while maternity is derived from behavioral observations and confirmed by genetic analyses (see ^73^ for detailed description).

### Pedigree data

To compare IBD-based estimates of realized relatedness to pedigree-based estimates, we applied the algorithm *TRACE* v0.1.0^77^ to pedigree data from 12,049 members of the Cayo Santiago rhesus macaque population, including 11,805 known mother-offspring dyads, 4,986 known father-offspring dyads, and up to 12 generations. For any given dyad (i.e., “focal” individuals), we calculated the relatedness coefficient (r_PED_) and categorized possible paths to a common ancestor into a kin class based on the depth of the path (i.e., the number of generations between the focals and their most recent common ancestor) and the sexes of the focals. Many dyads in our data set are related through multiple relatedness paths. However, as distant kinship contributes relatively little to IBD sharing^45^, we classified each dyad according to its primary kin class (i.e., closest consanguine relationship) for the present analysis. Thereby, we focused on the kin classes that are most commonly considered in studies on the Cayo Santiago macaques: parent-offspring, full siblings, grandparent-offspring, half siblings, half avuncular and half 1^st^ cousins^73^. Dyads that were related via more than one kin class of interest, as well as dyads that were not related via any of the kin classes of interest, were classified as “miscellaneous.” If a dyad shared no common ancestors in the pedigree, it was classified as nonkin (i.e., r_PED_ = 0).

### WGS data

We generated WGS data from 103 individuals who were selected as a representative subset of the pedigree. Specifically, we conducted a three-stage search, starting with a randomly chosen individual in the 12,049-individual Cayo Santiago pedigree. For this individual, we first selected all its first-degree relatives for whom DNA samples were available. In the second stage, we expanded the search to include all sampled, first-degree relatives of the individuals identified in the first stage.

Finally, we included all sampled, first-degree relatives of the individuals identified in the second stage. This iterative approach yielded a final sample of 103 individuals (57 females and 46 males) born between 1981 and 2010 (Tab. S1). Our sample included males with 1-24 and females with 1-8 offspring (Fig. 1B). We sequenced five of the 103 animals to high coverage (18.19-29.60×) for validating and optimizing our imputation and analysis pipeline. To complete the data set, we then sequenced 89 of the remaining individuals to 4-6× coverage. Finally, we included 9 individuals who had previously been sequenced to 6-38× coverage in a separate project (Tab. S1). For details on the DNA extraction and sequencing methods, see Supplementary Note 1.

### Analysis of WGS data

After quality control (removing bases with Phred-score of a base call < 20 starting from both 5’ and 3’ end of each read) and adapter trimming using *cutadapt*^78^, the resulting reads were aligned to the rhesus macaque reference genome *Mmul10*^79^ using *hisat2*^80^. We then removed likely PCR or optical duplicates using *Picard MarkDuplicates*.

### Benchmarking IBD segment calls

We used the five samples sequenced to high coverage (mean = 23.1×; Tab. S1) to produce a ground truth IBD set. We then performed downsampling experiments to investigate how IBD calling accuracy decays with decreasing coverage. First, to genotype the five high-coverage samples, we used *bcftools* v1.9^81^ to call genotypes at 23,874,572 predefined variant sites from the mGAP 2.4 reference panel^82^ with both minimum base call and minimum mapping quality set to 30. To remove low-quality genotypes, we retained only genotypes supported by between 15 and 100 reads and required that the genotype likelihood of the highest likelihood genotype was at least 1000 times greater than the second most likely genotype. Such filtering is important for calling high-quality ground truth IBD segments as most IBD callers are sensitive to genotyping errors.

After filtering, we used *IBIS* v1.20.9^83^ with default settings to call IBD for the five high-coverage samples. We chose *IBIS* for the high-coverage samples because it does not require phased data and performs well for high-quality diploid genotypes^83^. However, because *IBIS* is not designed for use with low-coverage WGS data, we used an alternative method, *ancIBD*, based on its superior performance compared to other IBD detection methods on low-coverage human ancient DNA data^67^. To assess the performance of *ancIBD* on imputed low-coverage data, we downsampled the five high-coverage samples to 6×, 4×, 2×,1×, 0.5× and 0.25× respectively, each with 25 independent replicates. We then used *GLIMPSE* 1.1^84^ to impute and phase the downsampled data using a haplotype reference panel of 741 Indian-origin *M. mulatta* individuals from mGAP 2.4^82^. We imputed genotype data for each downsampled replicate genome independently. We then explored various parameter combinations for *ancIBD* and chose a set that yields balanced performance for both precision and recall (see Tab. S2 for the list of parameters). We computed precision as the fraction of inferred IBD segments (within a specified length bin) that overlapped with an IBD segment in the ground truth data set of any length. We computed recall as the fraction of ground truth IBD segments (within a specified length bin) that overlapped an IBD segment of any length inferred in the downsampled data set. Finally, we used *ancIBD* with this modified parameter set to identify shared IBD segments between all individuals in the final data set. To avoid batch effects, we imputed all samples, including the high-coverage samples, before IBD calling.

### IBD2 calling

Many dyads in our complete sample (n_individuals_ = 103) are related via both maternal and paternal ancestors and consequently can have genomic regions where they share both alleles IBD. We designate such stretches as “IBD2” to distinguish them from segments where two individuals share only one haplotype IBD (“IBD1”). Excluding IBD2 in full siblings would lead to a considerable underestimation of relatedness, as they are expected to share a quarter of their genome in IBD2^10^. Hence, identifying IBD2 is crucial for obtaining an unbiased estimate of overall relatedness.

Therefore, we modified *ancIBD* to also identify IBD2 regions (Supplementary Note 2). We followed the same strategy as for IBD1 calling. Specifically, we first performed IBD2 calling in the five samples sequenced to high coverage using *IBIS*’s default cutoff of 2 cM to generate a ground truth data set. Next, we called IBD2 with the modifications applied to *ancIBD* in the 25 downsampled and imputed replicates of 6×, 4×, 2×,1×, 0.5× and 0.25× coverage respectively. We computed the sumIBD2_>2cM_ for each downsampled coverage level, averaged over 25 independent downsampled replicates. While the five high-coverage samples did not include any full siblings, they did include one parent-offspring trio (mother: 134814, father: 134854, son: 134834) in which the two parents are half 4^th^-cousins, which causes IBD2 sharing between parents and offspring. To ensure that IBD2 calling does not interfere with IBD1 calling in *ancIBD*, we compared the total sum of IBD1 regions and the number of IBD1 segments with and without calling IBD2. We found that the results are highly consistent, indicating little to no interference (Fig. S1).

### Processing of low-coverage samples

After the imputation of low-coverage data with *GLIMPSE*^84^, we performed several quality checks to identify and remove poorly imputed samples and variants. First, we computed average heterozygosity and the fraction of imputed sites for which the maximum genotype probability was greater than 99% (hereafter referred to as maxGP99 fraction; Tab. S1). The latter measure strongly correlates with coverage^67^. We retained samples with >90% maxGP99 and <35% heterozygosity. Out of 103 samples, 3 failed this quality control filter and were excluded from further analyses (Supplementary Note 3; Fig. S2). Second, we filtered out variants in the imputed data that violate Hardy-Weinberg-Equilibrium (HWE), using *plink* v1.90b3.29^39^ at a significance level of 0.01. Out of a total of 7,007,674 imputed variants with a minor allele frequency of at least 5%, 67,432 (0.96%) variants were excluded (Fig. S3). We found that removing these variants slightly improved the quality of called IBD segments, particularly by minimizing the rate at which single long segments were mistakenly subdivided into multiple shorter segments. Lastly, for IBD calling, we only used variants with a minor allele frequency of at least 5% (calculated from the reference panel), as imputation accuracy is generally much higher for common variants (Fig. S4)^84^.

### Global IBD sharing

To calculate the percentage of DNA shared between individuals, hereafter called r_IBD_, we summed the total length of shared IBD segments ≥ 8 cM per dyad (sumIBD1 + 2*sumIBD2) and divided it by the total diploid length of the macaque genome in Morgans (2 x 14.85 M = 29.70M). Here, we focused on segments ≥ 8 cM because longer segments offer clearer indications of recent ancestry^67^. Comparing r_PED_ and r_IBD_ revealed two cases of uncertain identity, which were excluded from further analysis (Supplementary Note 3; Fig. S5). Our final data set for IBD and realized relatedness analysis therefore included 98 individuals.

### Recombination map

As a recombination map for the *Mmul10* assembly^79^ was not available, we created a sex-averaged recombination map for the 20 rhesus macaque autosomes based on 25 unrelated Indian-origin individuals, randomly selected from the mGAP database v. 2.0^82^. After excluding indels, which can lead to phasing errors due to high genotyping error rates^85^, we first used *Beagle 5.2*^86^ to generate a set of phased macaque haplotypes based on SNP genotypes from all 2,054 rhesus macaques in the mGAP release 2.0^82^. We divided the autosomes into sections of 200k SNP markers, with an overlap of 20k SNPs between adjacent sections. These individual chunks were phased separately and then stitched together to create a final, phased VCF. We then used the software *LDhelmet* v1.10^87^ to compute a genetic map that estimates recombination in units of ρ/bp for the 25 animals. *LDhelmet* was run with the authors’ recommended settings: a window size of 50 SNPs, a block penalty of 5.0, a burn-in period of 10,000 iterations, and subsequent 1,000,000 iterations of the reversible-jump Markov Chain Monte Carlo algorithm. Subsequently, we identified and updated outlier regions with unusually high recombination rate estimates (or, unusually low levels of linkage disequilibrium) by setting the top 2.5% of ρ estimates to the value for the 97.5% quantile. Then, we calculated the cumulative ρ by multiplying the distance in bp between adjacent SNPs with the ρ/bp estimates and then summing these values across chromosomes to calculate the cumulative ρ. Next, we calculated a scaling factor for each chromosome by dividing the maximum cM value by the maximum cumulative ρ per chromosome^88^. By multiplying this scaling factor with each ρ estimate, we calculated the corresponding cM values^89^. Importantly, although linkage disequilibrium-based methods like *LDhelmet* are known to produce some errors^90–93^, the resulting recombination map is well-correlated with a previous map generated for *rheMac8*, an older version of the rhesus macaque reference genome (Fig. S6)^88^.

### Simulation of IBD segments

To validate our IBD results, we required a theoretical distribution of IBD segments and r_IBD_ for each kin class of interest (i.e. parent-offspring, full siblings, grandparent-offspring, half siblings, half avuncular, half 1^st^ cousins). To this end, we used *ped-sim* v1.4^94^ to simulate IBD segments for these kin classes, using the recombination map described above and a Poisson model for recombination. For each kin class, we ran 100 replicates. Since empirical IBD segments can only start or end with the first or last SNP per chromosome respectively, we truncated the simulated IBD segments so they did not extend beyond the positions where the first and last SNPs used in IBD inference occur.

### Estimating sex differences in recombination rate

Although constructing a fine-scaled sex-specific genetic map for rhesus macaques is beyond the scope of this study, our IBD estimates support a difference in recombination rates between females and males (see Results). We therefore designed a simple model to estimate the overall recombination rate ratio of females to males. Our model takes advantage of the difference in the number of IBD segments shared between maternally and paternally related grandparent-offspring and half sibling dyads, which must arise because of sex differences in recombination rates. Such differences have been observed in various human pedigrees^67,94^ and have been exploited to infer the sexes of ungenotyped ancestors^15^.

To estimate overall sex differences in recombination rate, we therefore focused on the number of shared IBD segments between grandparent-offspring and half siblings, considering only dyads that did not share any additional genetic relationship closer than 5^th^ degree. Let the male genetic map length (measured in Morgans) be 𝑟_𝑀_. If the female bias in recombination rate, in terms of foldchange increase relative to males is α, then the female genetic map length is 𝑟_𝐹_ = 𝛼𝑟_𝑀_. Denoting the number of meiosis between two individuals in a given kin class as 𝑑, and the number of autosomal chromosomes as 𝑐, then the expected number of IBD segments shared is proportional to 𝑟𝑑 + 𝑐 ^44^. In particular, the expected number of shared IBD segments for grandparent-offspring and half siblings in rhesus macaques, where the haploid autosome number is 20, are 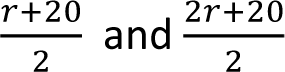 and respectively. Following Huff et al.^44^, we model the number of shared IBD segments using a Poisson distribution with mean 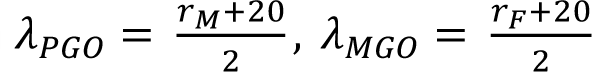 for paternal and maternal grandparent-2 offspring and 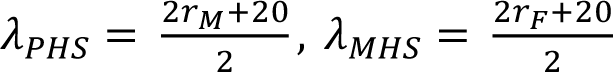 for paternal and maternal half siblings. Our likelihood model must therefore optimize over two free parameters: the genetic map length ratio α and the male genetic map length 𝑟_𝑀_. We sum up the Poisson log-likelihood for all dyads in these four kin classes, 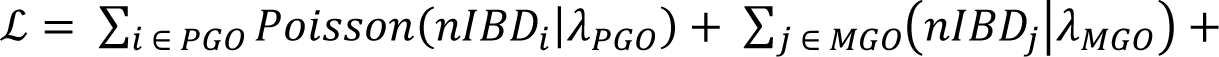 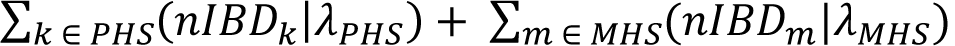. We then optimize the likelihood using the Nelder-Mead method in the *scipy.optimize.minimize* module with initial value 𝛼 = 1, 𝑟_𝑀_ = 14.85 (the sex-averaged genetic map length estimated in ^88^). To compute confidence intervals, we numerically calculate the Fisher Information of the likelihood function using the Python package *numdifftools*, and then approximate the 95% confidence interval by ± 1.96 × 𝑠𝑡𝑎𝑛𝑑𝑎𝑟𝑑 𝑒𝑟𝑟𝑜𝑟. We note that dyads of relatives are not fully independent because one individual can occur in multiple dyads of the same kin class. Because our likelihood function assumes independence, our confidence intervals are therefore likely to be underestimated.

In practice, we modified the values of λ as follows to account for the fact that (1) we are only able to detect IBD segments longer than 4 cM and (2) our findings revealed substantial background relatedness in the Cayo Santiago macaques (see Results), resulting in IBD even among individuals who have no known relatives in the pedigree. Non-negligible background relatedness in the sample likely arises from the small effective population size of this population and the small number of founders, followed by no subsequent immigration. To accommodate (1), we computed the percentage of the genome in IBD segments longer than 4 cM for simulated grandparent-offspring and half siblings (see Methods section *Simulation of IBD segments*). For (2), we computed 𝜆_𝑛𝑜𝑛𝑘𝑖𝑛,4𝑐𝑀_, the average number of IBD segments > 4 cM among nonkin dyads. Then, we updated the Poisson mean as follows, 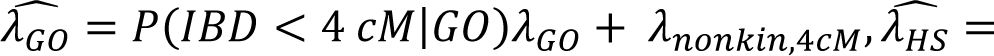 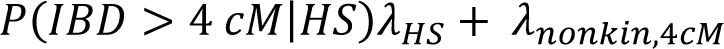. Note that this representation is not perfect, as the percentage of the genome in IBD segments longer than any given length threshold is expected to differ between maternally and paternally related dyads. However, this approximation should not introduce substantial bias: for instance, in humans, the percentage of the genome IBD segments > 4 cM differs by no more than 3% between maternally and paternally related grandparent-offspring and half sibling dyads (see Tab. S3 for simulation results).

We note that in theory the model only needs one free parameter, namely, α, because 𝑟_𝑀_ can be obtained directly from the known sex-averaged genetic map length: 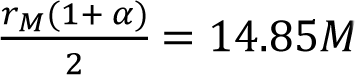. However, we found the model optimization based on α alone to produce unreliable results. Specifically, this approach led to higher estimated recombination rates in males than in females (i.e., α < 1) (𝛼^ = 0.990, 95% CI: 0.918-1.061), despite higher numbers of IBD segments in maternal compared to paternal dyads. In particular, this model predicts fewer IBD segments shared in maternal half-siblings (^𝜆^^ = 33.024 segments) than observed in our actual data (^𝜆^^ = 41.859). Consistent with this observation, comparing the two-parameter model against the one-parameter model using the Akaike Information Criterion (AIC) strongly favors fitting α and 𝑟_𝑀_ separately (delta AIC = 220.96), possibly because the two-parameter model better absorbs errors in IBD detection itself.

## Results

### Accuracy of IBD inference in low-coverage samples

Inferred IBD segments in downsampled data were highly consistent with our ground truth estimates from high-coverage data, even for coverage depths as low as 0.5 – 0.25x (Fig. 2). We identified four genomic regions (a total of ∼40 cM, ∼2.69% of the rhesus macaque genome) with very few SNPs, which made IBD calling problematic (Fig. S7). We thus masked these four regions (visualized as yellow blocks in Fig. 2, and we visualized one such example in Fig. S7) when calculating precision and recall. We note that these masked regions largely correspond to unresolved regions in the genetic map (Fig. 2, S6). Further, we found that some long segments in the empirical data are sometimes broken up into shorter ones due to sporadic genotyping errors, which is especially pronounced in parent-offspring dyads.

**Figure 2:**
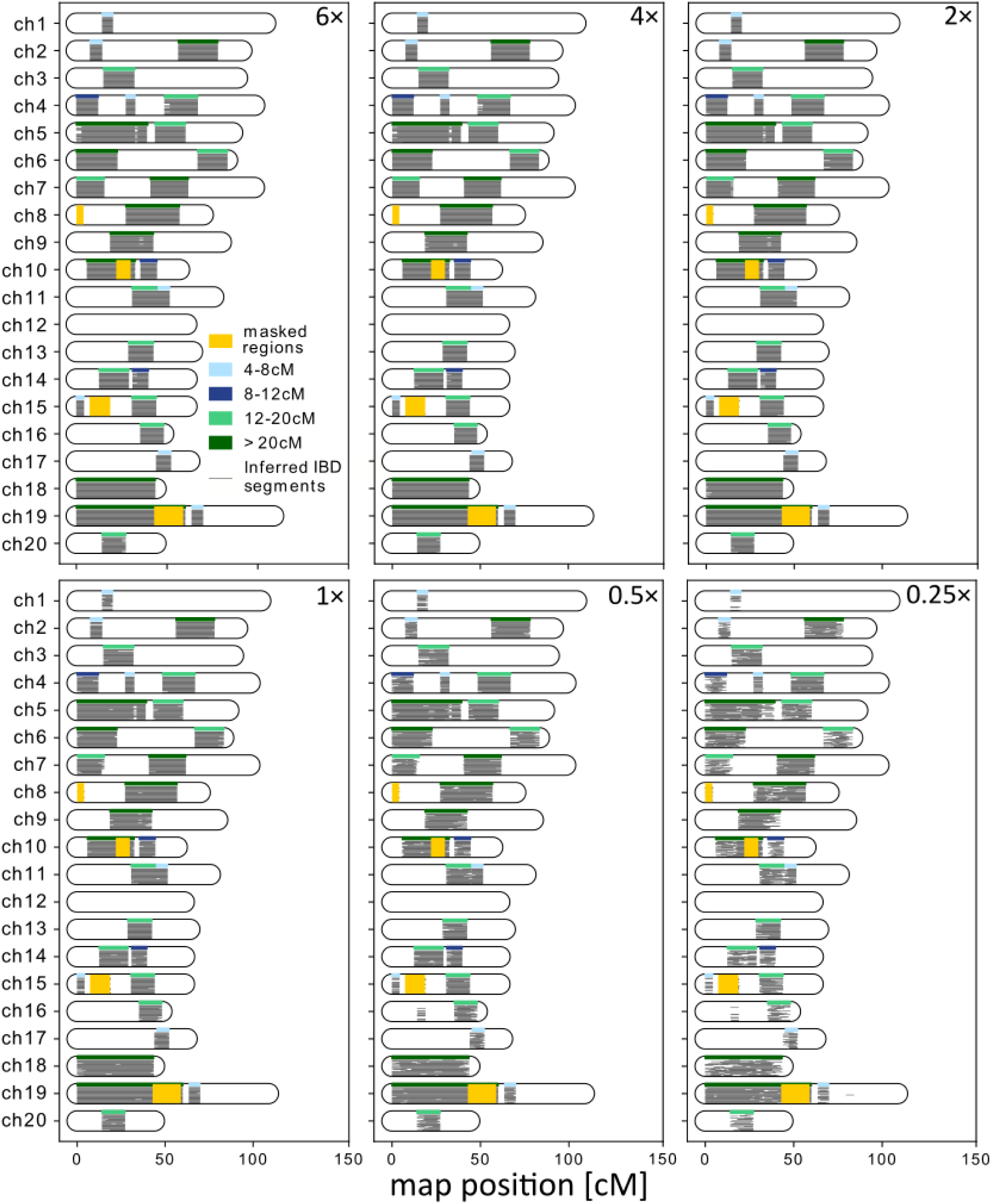
Example concordance in IBD calls from high-coverage and downsampled low-coverage data. Inferred and ground truth IBD1 for a niece and her half uncle (i.e., half brother of her parent, half avuncular). Ground truth IBD1 (called using IBIS on high-coverage data) are depicted as thick bars at the top of each chromosome and color-coded according to their length. IBD1 was inferred from downsampled data at 6×, 4×, 2×, 1×, 0.5× and 0.25× coverage. The 25 replicates are visualized as thin gray lines stacked together. Regions with few SNPs are masked with thick yellow bars. After imputation, we were able to reliably call long IBD segments even in data downsampled to 0.25× coverage.

We found that at coverages ≥ 0.5×, precision remains consistently high (>98%) for all length bins: [4 cM, 8 cM), [8 cM, 12 cM), [12 cM, 20 cM), and [20 cM, ∞), and recall is ≥80% even at 0.5× coverage (Fig. S8). At a coverage of 0.25× (Fig. S8), recall is less than 80% and IBD segments tend to be fragmented compared to those identified at higher coverages, resulting in a bias towards calling more, but shorter, IBD segments (Fig. 2). Consequently, while close relatives can still be identified at 0.25× from the total length of shared IBD, inferences that rely on the number and/or distribution of segment lengths (e.g. distinguishing maternal half siblings from paternal ones) will be prone to error.

Our analysis also showed that the performance of IBD2 calling in low coverage data was similarly robust (Fig. S9-S12). We computed the precision and recall for IBD2 in the same way as IBD1 but with different length bins: [2 cM, 4 cM), [4 cM, 8 cM), [8 cM, 12 cM), [12 cM, ∞) (Fig. S10). We found that precision is greater than 90% across all length ranges and all coverages tested here. Recall is poor for the 2-4 cM length bin and only reaches about 80% at 6×. But for > 4c M IBD2, recall is above 80% from for 1× or higher coverage. In addition, we calculated sumIBD2_>2cM_ for the two dyads used as ground truth IBD2 here (Fig. S11). The sumIBD2_>2cM_ converges to the ground truth value as coverage increases, consistent with the observation that our IBD2 calling has high precision at all coverages but low recall at low coverages. Finally, we computed sumIBD2_>2cM_ for all dyads of full siblings in the low-coverage data, and we found that the values fall around the expected value, which is one fourth of the total genome length (∼371.2 cM of 2970 cM; Fig. S12).

### IBD analysis highlights cryptic relatives and variance in realized relatedness

The final data set for IBD analysis included 98 individuals and 4,753 dyads: 93 parent-offspring dyads, 16 full siblings, 41 maternal and 39 paternal grandparent-offspring, 64 maternal and 469 paternal half siblings, 282 half avuncular dyads, and 384 half 1^st^ cousins. 438 dyads had an r_PED_ = 0 and were classified as nonkin. The remaining 2,927 dyads were either more distant relatives or followed more complex relatedness patterns and were classified as miscellaneous (Fig. 3, Tab. S4).

**Figure 3:**
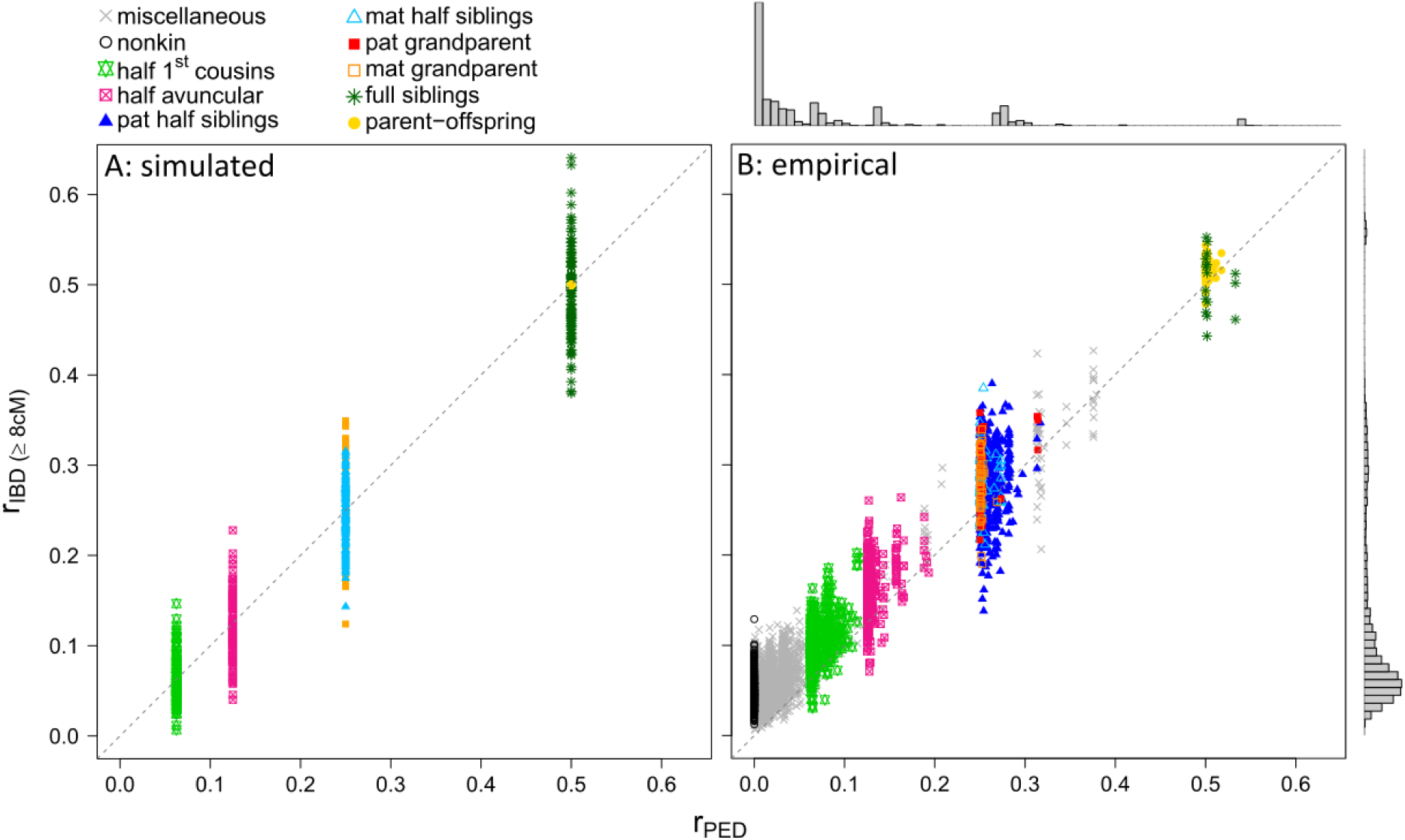
Comparison between IBD-based and pedigree-based kinship coefficients. The dashed line shows a perfect correlation between estimates. (A) shows simulated IBD estimates. As the simulation is based on a sex-averaged recombination map, we cannot distinguish between maternal and paternal grandparent-offspring (filled orange squares) nor half siblings (filled light blue triangles) here. (B) shows empirical IBD estimates. Based on the pedigree, dyads are color-coded according to their primary kin class (i.e., their closest consanguine relationship). Dyads of a given kin class that are shifted to the right on the x-axis relative to expectations for their primary kin class are related via more than one path (e.g. a half sibling dyad who is also half 2^nd^ cousins, produces an rPED value of 0.2656). Histograms show the distribution of rPED on top and rIBD on the right side for empirical data.

The simulated IBD segments provided a theoretical range of r_IBD_ for each kin class of interest. We found that the simulated and empirical IBD estimates for each kin class are highly concordant, such that the difference between the mean simulated and empirical r_IBD_ is less than 3% on average across all kin classes (Tab. 1, Fig. 3). r_PED_ and r_IBD_ are also highly concordant, with less than 3% difference in mean r_PED_ and r_IBD_ across all kin classes (Tab. 1, Fig. 3). However, within all non-parent-offspring dyads, r_IBD_ exhibits variation undetectable based on r_PED_ (Fig. 3). While some of this variation likely stems from estimation error, the variance in r_IBD_ for parent-offspring dyads – the only kin class where IBD sharing should be exact, in the absence of inbreeding – is the lowest among all the kin classes we considered. This result, in addition to the high accuracy of IBD estimation in downsampled data (> 90% precision and recall for >= 1× coverage), suggests that our IBD-based pipeline provides information about realized relatedness beyond that contained in the pedigree alone.

**Table 1:**
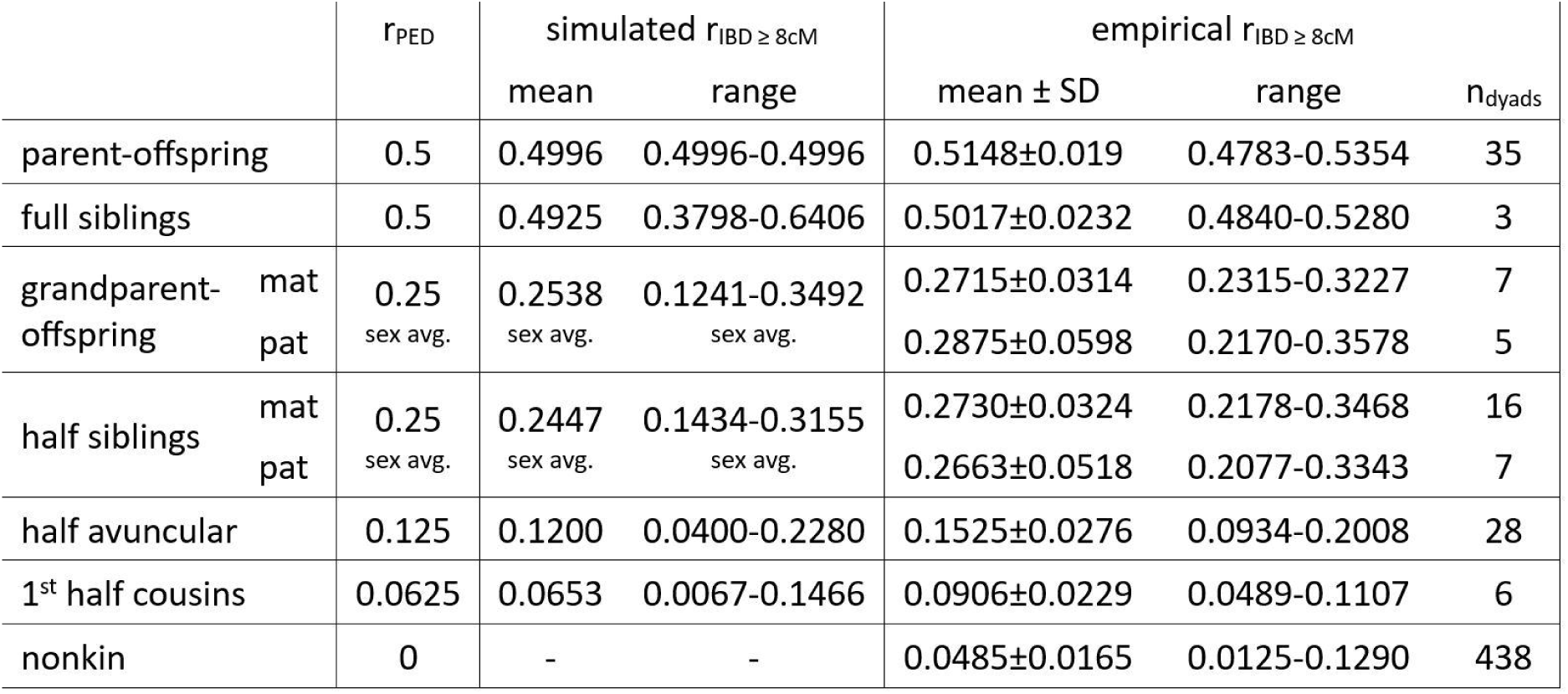
Mean and range of rPED and rIBD in the simulated and empirical data set for each kin class of interest (nindividuals = 98). Numbers based on the empirical data set only include dyads that do not share any additional distant kin relationships. Simulations are based on a sex-averaged recombination map, so we did not distinguish IBD estimates based on maternal versus paternal kin relationships. Dashes indicate values that were not assessed.

Interestingly, our empirical IBD estimates tended to be biased upwards relative to expectations from the pedigree (mean r_IBD_ = 0.110, median r_IBD_ = 0.0659 compared to mean r_PED_ = 0.076, median r_PED_ = 0.022; Tab. 1; Fig. 3B, S13). Half avuncular and half 1^st^ cousins share higher numbers of segments and sums of IBD1 than expected based on the simulation (Fig. 4). Further, individuals with no known shared ancestors in the pedigree data (i.e., dyads classified as nonkin) also shared an estimated 4.85% ± 1.65% SD of their genomes in large IBD segments, compared to pedigree-based expectation of 0 (Tab. 1, Fig. 5G, S13). Combined, these observations suggest substantial background relatedness in the Cayo Santiago population. Such a result could arise due to relatedness in the population founders (assumed to be 0 in pedigree-based estimates) or incomplete pedigree data. Our analysis suggests contributions from both. Specifically, when excluding all individuals with at least one parental ID missing (n_individuals_ = 4, n_dyads_ = 142) or all individuals with at least one grandparental ID missing (n_individuals_ = 34, n_dyads_ = 374), the median r_IBD ≥ 8cM_ of nonkin dropped to 0.0454 and 0.0405, respectively, but still did not approach 0 (Fig. S13). In this data set, seven nonkin dyads with missing parental and/or grandparental ID have an r_IBD ≥ 8cM_ = 0.09 - 0.13, which corresponds to a 3^rd^ (i.e., half avuncular) or 4^th^ (i.e., half 1^st^ cousins) degree relationship in our study population (Tab. 1).

**Figure 4:**
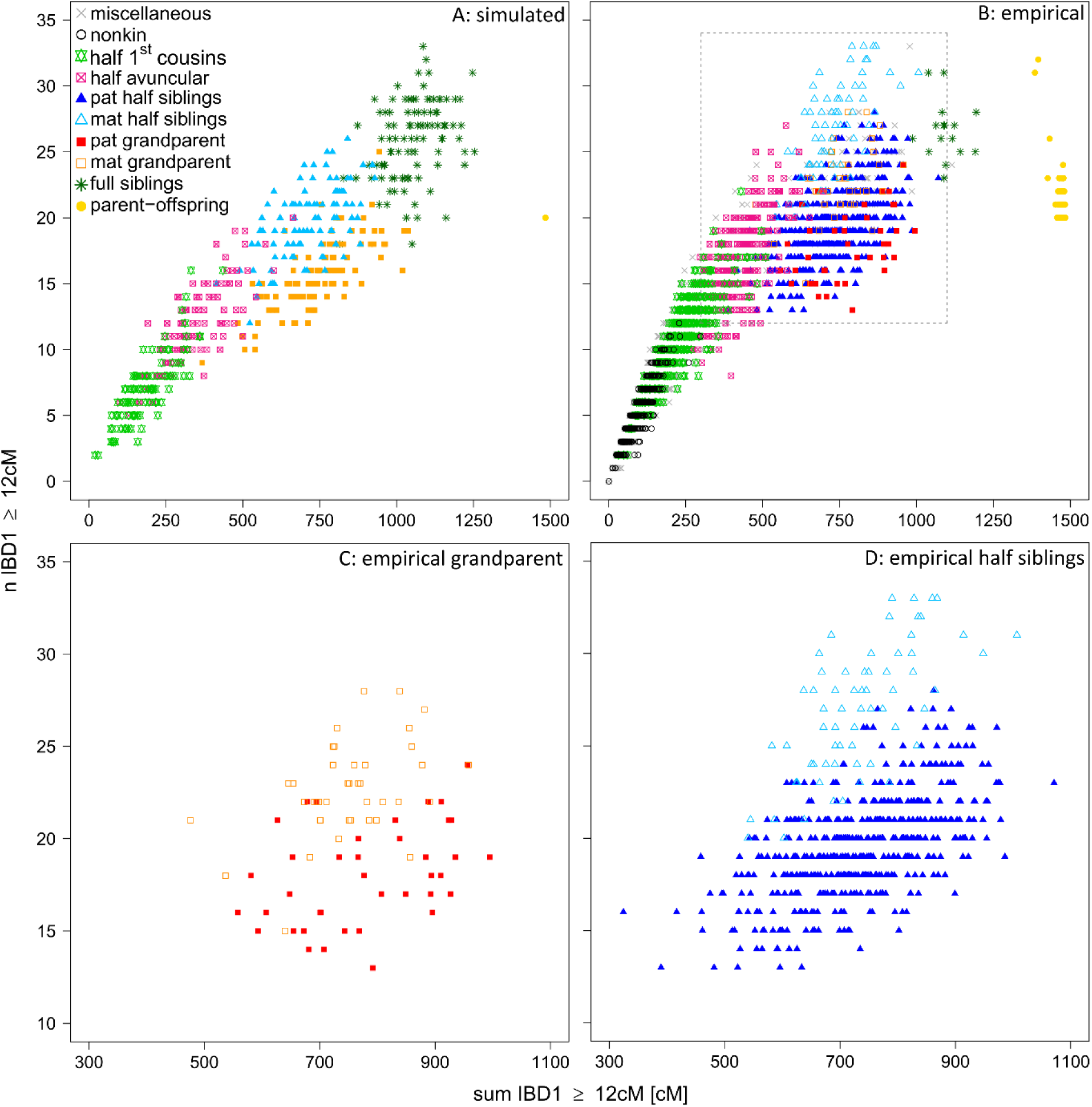
Distribution of number and length of shared IBD1 segments. Based on the pedigree, dyads are color-coded according to their primary kin class (i.e. their closest consanguine relationship). Gray crosses are dyads with more distant and/or more complex relatedness patterns. (A) shows simulated data. As the simulation is based on a sex-averaged recombination map, we cannot distinguish between maternal and paternal grandparent-offspring (filled orange squares) or half siblings (filled light blue triangles). (B) shows empirical data. The gray dashed frame shows the area expanded in (C) for grandparent-offspring and (D) half sibling relationships, where the difference between maternally and paternally related dyads is visually apparent.

**Figure 5:**
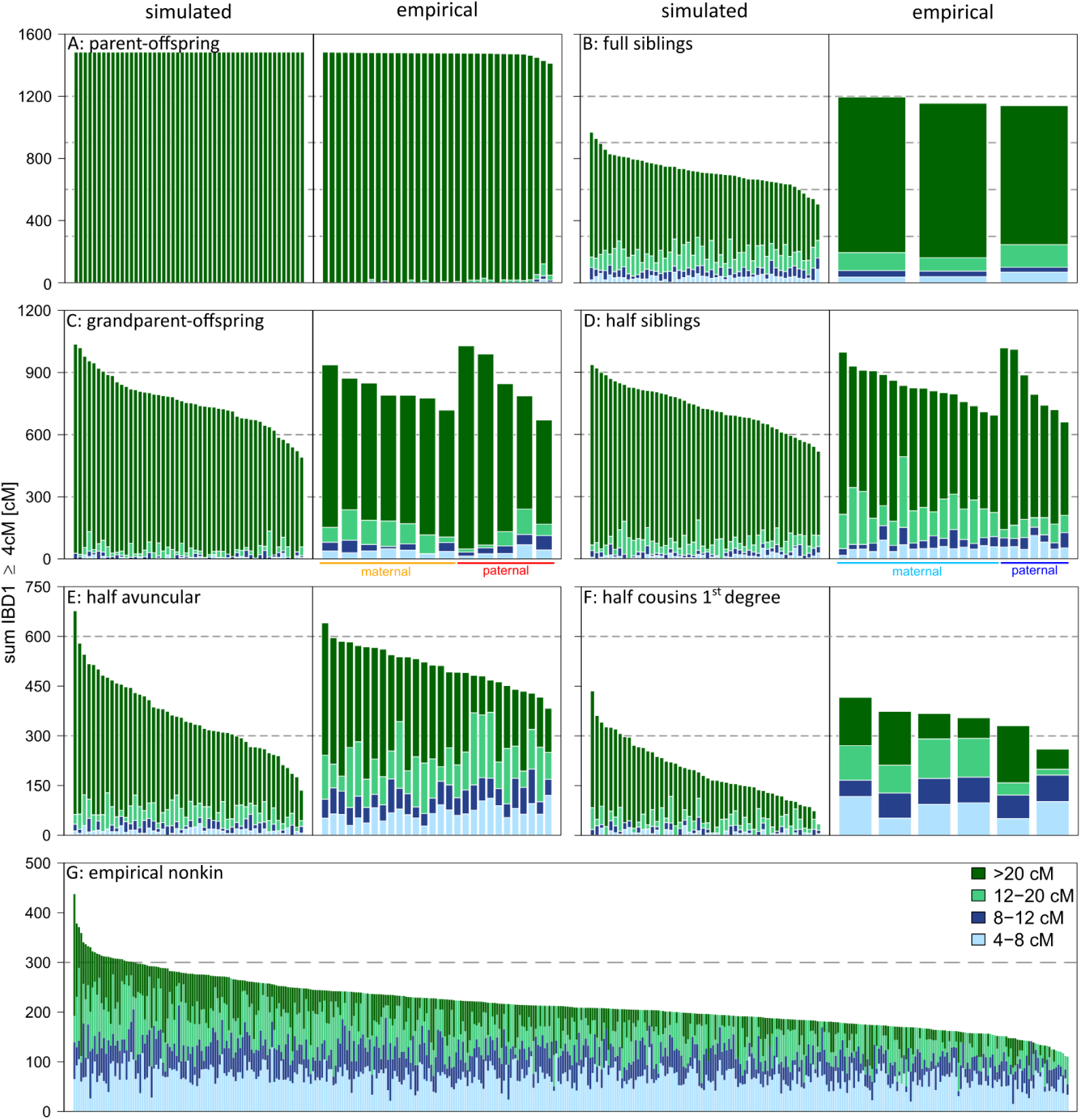
Size class of shared IBD segments. (A) - (F) The left panels show simulated data and the right panels show corresponding empirical data. (G) shows unrelated dyads with rPED = 0. Each bar represents one dyad, with the dyads sorted from the largest to the smallest total sum of shared IBD. The size classes are color-coded and vertically ordered so that the longest segments are shown on top of each bar and the smallest on the bottom. The simulated data are not sex-specific and display only 50 out of 100 simulations. The empirical data set only includes dyads that do not share any additional distant kin relationships. In the empirical data, we distinguish between maternal and paternal half siblings.

### Sex differences in recombination rate produce distinct patterns of IBD in maternal versus paternal relatives

To estimate differences in recombination rate between males and females, we used data from 41 maternal grandparent-offspring, 35 paternal grandparent-offspring, 64 maternal half siblings and 425 paternal half siblings. We estimate that the overall recombination rate is ∼1.502x higher in females than males (95% CI: 1.406 - 1.598; Tab. S3). Our estimated male genetic map length is 16.789 M (95% CI: 16.251 - 17.326), which is substantially longer than the previous estimate of 14.85 M^88^. This discrepancy might be due to 1) IBD detection errors, for example, occasional false positives and the break-up of long, continuous IBD segments into several adjacent shorter ones; 2) Xue et al.^88^ based their estimate on an older assembly than the *Mmul10* assembly used here, which is less complete. However, we note that the discrepancy of estimated genetic length is unlikely to substantially bias the estimated female-to-male recombination rate ratio because the IBD detection error rates are expected to be similar across maternal and paternal dyads and therefore the relative ratio of the number of segments of maternal dyads to that of paternal dyads is expected to remain the same. More precisely, denote 𝜆_𝑀_, 𝜆_𝑃_ as the ground truth number of IBD segments in maternal and paternal dyad, respectively. We model the IBD detection error as follows: each dyad on average has 𝑛 false positives, and each ground truth segment has probability 𝑝 of being inferred as two adjacent shorter segments. Then the number of detected IBD segments can be written as (1 + 𝑝)𝜆_𝑀_ + 𝑛, (1 + 𝑝)𝜆_𝑃_ + 𝑛. Since 𝑛 is significantly smaller than 𝜆_𝑀_, 𝜆_𝑃_ (because our detection algorithm has high precision), the ratio 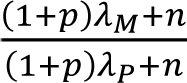 is approximately 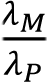. The sex differences in recombination rate lead to differences in IBD sharing between maternal and paternal relatives. To compare the number and cumulative length empirical IBD1 segments, we focused on segments ≥ 12 cM, which should reflect recent shared ancestry rather than background relatedness. We observe that segments are split up into several shorter ones in maternal grandparent-offspring and half siblings compared to paternal grandparent-offspring and half siblings, while the total IBD1 sum is similar (mean sumIBD1 ≤ 12 cM: maternal grandparent-offspring 165 cM, paternal grandparent-offspring 131 cM, maternal half siblings 269 cM, paternal half siblings 181 cM; Fig. 4C, 4D, 5C, 5D).

## Discussion

Here, we benchmarked and applied IBD-based methods to low-coverage sequencing data from a free-ranging population of rhesus macaques. By applying and adapting methods originally tailored to ancient DNA data from humans, we demonstrate the possibility of robustly calling IBD segments in nonhuman primates. The resulting estimates allow us to measure the biological gradient in realized relatedness within kin categories, opening the door to substantially increased resolution into the role of kinship in the evolution of sociality and the consequences of inbreeding.

For wild animals, DNA samples of adequate quantity and quality may not be available to produce genotype data of sufficient quality to apply common IBD calling methods. Additionally, producing WGS data at high coverage (≥ 20×) remains cost-prohibitive for large numbers of individuals. An alternative cost-effective approach is to produce WGS data at low coverage (< 10×) and impute diploid genotypes based on a high-quality reference genome panel^66,95^. In this study, we produced high-coverage data for a small subsample of individuals, to establish and test an imputation pipeline (Fig. 1C). We found that, for our study population imputed against the mGAP reference panel, IBD segments ≥ 4 cM can be robustly detected even for 0.5× WGS data (Fig. 2, S8). Imputation is therefore a very promising approach for future genome-wide analyses, though the performance may vary depending on the genetic diversity of the studied population (i.e. if there are enough SNPs available to infer IBD segments), the accessibility of DNA samples of sufficient quality and quantity, and the availability of a genomic reference panel. Given that reference panels are becoming increasingly possible to develop for non-model organisms^66^, we anticipate that methods like the one we develop here will soon become applicable in many species.

Our empirical distribution of IBD segments largely mirrors the simulated data, showing that measurement errors are small compared to true biological signal. We observed that the variance in distant kin is very similar between empirical and simulated data (Tab. 1, Fig. 3, 4). It is noteworthy that the range of r_IBD_ in empirical full siblings is considerably smaller than predicted by the simulations, which is evidence for crossover interference in rhesus macaques. Caballero et al.^94^ showed in humans that models including crossover interference perform better in estimating empirical patterns of IBD sharing. However, since no recombination interference model is available for rhesus macaques thus far, we ran the simulations using a Poisson model, which produces good approximations for distant kin, but less precise predictions of IBD sharing patterns in close kin. Thus, comparisons between simulated and empirical IBD sharing patterns in full siblings may need to be treated with caution. We further observed that empirical parent-offspring share more IBD segments than the number of autosomal chromosomes. This result occurs because true IBD segments get split up into several smaller segments (Fig. 4, 5), because of genotyping errors introduced in imputation. However, parent-offspring can still clearly be distinguished from all other kin classes (Fig. 4).

Compared to commonly used methods to estimate kinship (e.g. pedigrees, STR markers), IBD-based methods provide more insight into relatedness structures that are potentially relevant to answering biological questions. First, IBD-based methods can capture the substantial biological variation of kin classes, such that r_IBD_ faithfully represents the gradient of realized relatedness within classes. The possibility to precisely measure this gradient opens new avenues, for example, in the study of kin discrimination in behavioral ecology. Individuals may prefer one sibling over another within the same pedigree kin class due to differences in realized relatedness. On the other hand, individuals may actively avoid mating with kin sharing higher realized relatedness to prevent fitness costs due to inbreeding. Further, the variation of realized relatedness within kin classes leads to overlapping relatedness values across kin classes. For example, although individuals share more DNA with their half siblings than their half avunculars on average, for ∼17% of individuals in our data, this pattern can be reversed (Fig. 3, Tab. S4). This raises the question of whether traditional kin classes used in behavioral studies reflect kinship as perceived by the animals themselves.

Second, IBD-based methods can help to resolve cryptic genetic relationships, which is particularly interesting for field studies with gaps in the pedigree. In the current analysis, nonkin dyads nevertheless share long segments of their genomes IBD. Our finding that IBD sharing among nonkin decreases – although does not disappear – when individuals missing a parental and/or grandparental ID in the pedigree are excluded (Fig. S13) suggests that these dyads share common ancestry that cannot be captured using r_PED_. For example, in our study, the IBD sharing patterns of seven nonkin dyads with especially high r_IBD_ are consistent with half avuncular or half 1^st^ cousins, indicating that those dyads share a 3^rd^ or 4^th^ degree relatedness through the missing parent (Tab. S4). While even a few such gaps in a pedigree can lead to inaccurate estimates^17,19,20^, IBD-based approaches can detect such genetic relationships. Doing so is likely to be particularly important in quantitative genetic studies, in which accurate measures of genetic relatedness are required for accurate heritability estimation, and in association analyses, to appropriately correct for population structure^6^.

Third, we found that our study population generally shares more IBD than predicted by the simulations and the pedigree, which is especially visible in parent-offspring dyads, where a considerable fraction of the genome is IBD for both alleles (i.e., IBD2; Tab. 1, Fig. 3). This pattern indicates that the parents are minimally distant relatives, in line with the observation that all dyads in our sample share some IBD segments (Tab. 1, Fig. 5G). This background relatedness could arise from several causes, including a relatively small founder population that may have included related individuals, a well-documented population bottleneck in 1972, the lack of genetic influx since the population founding, and strong reproductive skew in males, all of which can lead to higher IBD sharing^53,71,96^. Estimates of background relatedness therefore provide information about the demographic history and the effective population size of a study population, which also have ramifications for conservation and population viability^63,64^.

Fourth, IBD-based methods are sensitive to sex differences in recombination rate, which, in eutherian mammals, tend to be higher in females than males^48^. Not only do they allow these differences to be estimated – here, we inferred a recombination rate ∼1.502x higher in females than males, in line with an earlier estimate of 1.513x reported in Coop and Przeworski^97^ – they also suggest a strategy for reconstructing if a dyad (e.g., grandparent-offspring or half-siblings) is related via shared female or shared male ancestors. For example, our current analysis shows that, although maternal half-sibs and paternal half-sibs share the same cumulative amount of the genome IBD (on average), the proportion of the genome IBD is broken up across more segments in maternal half-sibs than paternal half-sibs. This information is important in many settings, including understanding the mechanisms underlying inbreeding avoidance and kin-biased behavior, estimating maternal effects on trait variation, and investigating the role of vertical transmission.

To conclude, here we applied and tested IBD methods to estimate realized relatedness in a free-ranging population of nonhuman primates. The resulting approach is applicable for IBD calling in a wide array of species and populations, which makes it possible to estimate relatedness in populations for which no pedigree data are available. Our results demonstrate that robust IBD calling is possible even at coverage as low as 0.5×. Further, we found that IBD-based methods capture considerable variation in relatedness beyond what can be inferred from pedigrees alone. The possibility to robustly measure realized relatedness provides a novel tool to understand the role of biological relatedness in a wide range of disciplines.

## Code availability

Code for analyzing the high-coverage sequencing data, comparing downsampled data sets followed by imputation, and imputation and IBD calling in low-coverage data is available on GitHub: https://github.com/vmjovanovic/macaqueIBD

## Data availability

Sequencing data for one sample are already available on NCBI Sequencing Read Archive (SRA: https://www.ncbi.nlm.nih.gov/sra/; see Tab. S1 for accession code). We will make the rest of the aligned sequencing data included in the final data set publicly available via NCBI SRA, once the manuscript is accepted for publication.

## Author contributions

Our annotation of author contributions follows the CRediT Taxonomy labels (https://casrai.org/credit/). In instances where multiple authors fulfil the same role, their degree of contribution is identified as either ’lead,’ ’equal,’ or ’support’. **Conceptualization** - lead: AW; support: NSM, JT, KN. **Data curation** - equal: AF, VMJ, SB, ARL, AW. **Formal analysis** - lead: AF, VMJ, YH, DFC; support: HR, NSM, BM, JT. **Funding acquisition** - lead: AW, support: NSM, DFC, LJNB, MLP. **Methodology** - equal: AF, VJM, YH, HR. **Project administration** - equal: AF, AW. **Resources** - equal: NSM, MJM, JEH, LJNB, MLP, ARL, KN, HR, AW. **Software** - equal: YH, HW, HR; support: PFS. **Supervision** - lead: AW; support: KN, HR. **Visualization** - lead: AF; support: YH. **Writing (original draft)** - lead: AF; support: VMJ, YH, HR, AW. **Writing (review & editing)** - equal: AF, VMJ, YH, NSM, DFC, BM, MJM, HW, PFS, SB, JEH, LJNB, MLP, ARL, JT, KN, HR, AW.

## Conflict of interest

The authors declare no conflict of interest.

## Acknowledgements

We thank the Caribbean Primate Research Center (CPRC), especially Melween Martinez, Carlos A. Sariol Curbelo and all field staff, for their continuing support of our work. We thank Richard McElreath for providing data storage and for his thoughtful comments on this manuscript. We thank Peter Fröhlich for IT support. We are grateful to Rebecca Bellone and Robert A. Grahn for providing DNA samples from the University of California-Davis sample repository. We are thankful to Antonia Krüger for managing the DNA samples and preparing them for sequencing. We thank the Cologne Center for Genomics (CCG), especially Kerstin Becker and Janine Altmüller, for producing the sequencing data. We are grateful to Connor Walen, Mitchell Sanchez Rosado and James Higham, who supervised the shipment of DNA samples. Thomas Gatter and Lars Kulik provided great support for calculations of the pedigree relatedness. Marina Watowich gave us valuable advice for imputation. Brigitte M. Weiß supported us in planning the project. The vast majority of sequencing costs as well as graduate funding was provided by the Deutsche Forschungsgemeinschaft (DFG) within a sequencing initiative (grant number WI 1808/7-1, project number 433163659 approved to AW). Funding for the additional nine individuals used from previous studies was provided by the National Institutes of Health (NIH; R01-MH-096875 to MLP; R01-MH-118203 to MLP, LJNB, NSM; R01-AG-060931 to LJNB, NSM). Cayo Santiago is supported by the Office of Research Infrastructure Programs (ORIP) of the NIH (P40 OD012217). DFC is supported by NIH grants U24HG012483 and U24MH123696 (both to DFC) and NIH Office of Directors (NIH/OD) grant P51OD011092 (to the Oregon National Primate Research Center). The content of this publication is solely the responsibility of the authors and does not necessarily represent the official views of NIH or ORIP.

## Notes

### Competing Interest Statement

The authors have declared no competing interest.

