## Supplementary Material for "Taking identity-by-descent analysis into the wild: Estimating realized relatedness in free-ranging macaques"

[1]Behavioral Ecology Research Group, Faculty of Life Sciences, Institute of Biology, Leipzig University, Leipzig, Germany; [2]Department of Primate Behavior and Evolution, Max Planck Institute for Evolutionary Anthropology, Leipzig, Germany; [3]Human Biology and Primate Evolution, Institut für Zoologie, Freie Universität Berlin, Berlin, Germany; [4]Bioinformatics Solution Center, Freie Universität Berlin, Berlin, Germany; [5]Department of Archaeogenetics, Max Planck Institute for Evolutionary Anthropology, Leipzig, Germany; [6]Bioinformatics Group, Institute of Computer Science, and Interdisciplinary Center for Bioinformatics, Leipzig University, Leipzig, Germany; [7]Center for Evolution & Medicine, School of Life Sciences, Arizona State University, Tempe, USA; [8]Division of Genetics, Oregon National Primate Research Center, Portland, Oregon, USA; [9]Department of Neuroscience, Perelman School of Medicine, University of Pennsylvania, Philadelphia, PA, USA; [10]Max Planck Institute for Mathematics in the Sciences, Leipzig, Germany; [11]Institute for Theoretical Chemistry, University of Vienna, Austria; [12]Facultad de Ciencias, Universidad Nacional de Colombia, Bogotá, Colombia; [13]Santa Fe Institute, Santa Fe, NM, USA; [14]Department of Biological and Biomedical Sciences, North Carolina Central University, North Carolina, Durham, USA; [15]Research and Collections Section, North Carolina Museum of Natural Sciences, North Carolina, Raleigh, USA; [16]Department of Biological Sciences, North Carolina State University, North Carolina, Raleigh, USA; [17]Department of Evolutionary Anthropology, Duke University, North Carolina, Durham, USA; [18]Renaissance Computing Institute, University of North Carolina at Chapel Hill, Chapel Hill, North Carolina, USA; [19]Centre for Research in Animal Behaviour, University of Exeter, Exeter, UK; [20]Marketing Department, the Wharton School of Business, University of Pennsylvania, Philadelphia, PA, USA; [21]Department of Psychology, School of Arts and Sciences, University of Pennsylvania, Philadelphia, PA, USA; [22]Cayo Santiago Field Station, Caribbean Primate Research Center, University of Puerto Rico, Punta Santiago, Puerto Rico; [23]Department of Biology, Duke University, Durham, North Carolina, USA; [24]Duke University Population Research Institute, Durham, North Carolina, USA; [25]German Centre for Integrative Biodiversity Research (iDiv), Halle-Jena-Leipzig, Germany

*AF, VMJ and YH contributed equally as first authors

#KN, HR and AW contributed equally as senior authors

January 2024

### Supplementary Note 1: DNA extraction and whole genome sequencing

Nine out of 103 samples were sequenced in a parallel project (for DNA extraction and sequencing methods see [1]). The remaining 94 samples were processed as follows: At Leipzig University and Arizona State University, DNA was extracted from whole blood samples using the DNeasy® Blood Tissue Kit (Qiagen). At the University of California-Davis, DNA was extracted from white blood cells following their separation from whole blood samples using centrifugation. Specifically, 75 µl of the isolated white blood cells were mixed with 1.2 mL of a 10 mM NaCl-10 mM EDTA solution at pH 7.0 to eliminate any remaining serum and red blood cells. The samples were then subjected to centrifugation at 16,000g for 1 minute, and the supernatant was removed. 100 µl of 200 mM NaOH was added to each tube containing the pelleted white blood cells. The tube was agitated to resuspend the pellet and then incubated for 15 minutes at 97°C. After incubation, each sample was treated with 100 µl of 200 mM HCl-100 mM Tris-HCl at pH 8.5.

The quantity of DNA in the samples was estimated by using a Qubit Fluorometer (Life Technologies). All samples were diluted to obtain a DNA quantity of either 600 ng, 300 ng, or 25 ng per sample. In three cases, the extraction yielded less than 25 ng of DNA (Tab. S1) and these samples were processed differently for WGS (see below).

Libraries were prepared with either the Nextera DNA (Illumina) or DNA Tagmentation (Illumina) preparation kit. We followed a PCR-free protocol for samples containing > 25 ng of DNA, while samples containing < 25 ng were processed with 12 PCR cycles. Library preparation was followed by clean up and/or size selection using SPRI beads (Beckman Coulter Genomics). After library quantification (Qubit, Life Technologies) and validation (Agilent TapeStation, only for libraries generated using PCR), equimolar amounts per library were pooled. The pools were quantified using the Peqlab KAPA Library Quantification Kit and the Applied Biosystems 7900HT Sequence Detection System. The library pools were sequenced on an Illumina NovaSeq6000 sequencing instrument at the Cologne Centre for Genomics, Germany using paired-end 2×150 bp reads.

### Supplementary Note 2: Extension of *ancIBD* to call IBD2

To call IBD2, we extended the functions of *ancIBD*[2] as follows. Denote the two haplotypes of the first sample as (1A, 1B) and that of the second sample as (2A, 2B). The default *ancIBD* hidden-Markov model (HMM) has five hidden states, namely, a non-IBD state and four IBD states (1A/2A, 1A/2B,

1B/2A, 1B/2B). Each of the four IBD states encodes one of the four possible configurations for two diploid samples to share identical haplotypes. 1A/2A, for example, denotes the case where the haplotype A of sample 1 is identical to the haplotype A of sample 2. To detect IBD2 regions, we added two more hidden states, (1A/2A & 1B/2B) and (1A/2B & 1B/2A). These two IBD2 states each encode one of the two possible configurations for two diploid samples to have identical haplotypes. (1A/2A & 1B/2B), for example, denotes the case where haplotype A of sample 1 is identical to haplotype B of sample 2 and haplotype B of sample A is identical to haplotype B of sample 2. To account for the additional two hidden states, we also extended the transition matrix as follows,

$$\begin{array}{l} nonIBD \\ IBD1_1 \\ IBD1_2 \\ IBD1_3 \\ IBD1_4 \\ IBD2_1 \\ IBD2_2 \end{array} \begin{bmatrix} & \frac{\sigma_1}{4} & \frac{\sigma_1}{4} & \frac{\sigma_1}{4} & \frac{\sigma_1}{4} & 0 & 0 \\ \sigma_2 & & \frac{\sigma_3}{4} & \frac{\sigma_3}{4} & \frac{\sigma_3}{4} & \frac{\sigma_1}{2} & \frac{\sigma_1}{2} \\ \sigma_2 & \frac{\sigma_3}{4} & & \frac{\sigma_3}{4} & \frac{\sigma_3}{4} & \frac{\sigma_1}{2} & \frac{\sigma_1}{2} \\ \sigma_2 & \frac{\sigma_3}{4} & \frac{\sigma_3}{4} & & \frac{\sigma_3}{4} & \frac{\sigma_1}{2} & \frac{\sigma_1}{2} \\ \sigma_2 & \frac{\sigma_3}{4} & \frac{\sigma_3}{4} & \frac{\sigma_3}{4} & & \frac{\sigma_1}{2} & \frac{\sigma_1}{2} \\ 0 & \frac{\sigma_1}{4} & \frac{\sigma_1}{4} & \frac{\sigma_1}{4} & \frac{\sigma_1}{4} & & \frac{\sigma_3}{2} \\ 0 & \frac{\sigma_1}{4} & \frac{\sigma_1}{4} & \frac{\sigma_1}{4} & \frac{\sigma_1}{4} & \frac{\sigma_3}{2} & \end{bmatrix}$$

Where $\sigma_1$ is the transition rate from nonIBD to IBD1 state, $\sigma_2$ is the transition rate from IBD1 state to nonIBD, and $\sigma_3$ is the transition rate within the four IBD1 states and the two IBD2 states. The transition from IBD2 state to nonIBD state (and vice versa) is virtually impossible because it requires two recombination events (one in the maternal germline and one in the paternal germline) to occur at the same genomic locus. Therefore, we set the rates for transitions from the two IBD2 states to the nonIBD state to zero. Parameters used for IBD calling are listed in Tab. S2. To calculate the emission probability for the IBD2 state, we follow the notation in Supplementary Note 1 from Ringbauer et al. [2]. We denote the emission probability for the case (1A/2A & 1B/2B) as the 6$^{th}$ hidden state (s = 6). We use the same notation as in Ringbauer et al.[2]. Throughout, we denote reference and alternative alleles as 0 and 1, respectively, and the corresponding genotype as $g \in \{0,1\}$. The observed data $D$ of our emission model will be the haploid dosage, which is the probability of a phased haplotype carrying an alternative allele, here denoted for each haplotype $h$ as $x_h = P(g_h = 1),\ h \in \{1A, 1B, 2A, 2B\}$. Further, we denote the phased genotypes of two diploid samples as $g = (g_{1A}, g_{1B}, g_{2A}, g_{2B})$, and $p$ be the allele frequency for the derived allele, then following the components listed in Tab. S5, the emission probability for $s = 6$ can be calculated as follows,

$$P(D|s=6) \sim \sum_{g \in \mathcal{G}} \frac{P(g|D)}{P(g)} P(g|s=6) = \frac{(1-x_{1A})(1-x_{1B})(1-x_{2A})(1-x_{2B})}{(1-p)^2} + \frac{x_{1A}(1-x_{1B})x_{2A}(1-x_{2B})}{p(1-p)} +$$

$\frac{(1-x_{1A})x_{1B}(1-x_{2A})x_{2B}}{p(1-p)} + \frac{x_{1A}x_{1B}x_{2A}x_{2B}}{p^2}$. The case for (1A/2B & 1B/2A) can be derived analogously and is omitted here.

### Supplementary Note 3: Excluded samples

Out of 103 samples, three samples did not pass quality filters: Two samples (193096, 193079) had very low maxGP99 fraction (< 80%). Further investigation revealed high amounts of bacterial contamination (running reads through BLAST, Madden 2013) and only low amounts of macaque DNA (i.e. coverage of 0.015× and 0.22×, respectively). One sample (163713) had very high average heterozygosity, likely due to cross-contamination. These three samples were excluded from further analysis. Furthermore, comparing $r_{PED}$ and $r_{IBD}$ revealed two individuals with uncertain identity: $r_{IBD}$ between these individuals and their putative close and distant kin strongly mismatched $r_{PED}$ (Fig. S4). Therefore, we concluded that the DNA must originate from different individuals of yet unknown identity and consequently, these samples were excluded from further analysis. This resulted in a final sample size of 98 samples.

### Tables and Figures

**Table S1:** Information about sex, birth season, coverage, heterozygosity and maxGP99 fraction for each sample that was sequenced in this study. Gray shaded rows indicate individuals that are not included in the final data set. The samples were excluded because the value marked in red lay outside of the tolerated range (see *Processing of low-coverage samples* for details). For individuals of uncertain identity, we have marked sex and birth season as NA, as we do not know the true identity of these individuals (see *Global IBD sharing* for details).

**Table S2:** *ancIBD* parameters used in this study. The parameters used in this study are different from the default parameters of *ancIBD*, which is fine-tuned for human ancient DNA data. Here, we chose parameters to have a balanced performance on precision and recall of both IBD1 and IBD2 segments. For parameters not listed in this table, we used their default values. We note that we have used a very lenient value for the posterior probability cutoff for IBD2 segments. We have found this to substantially boost power while introducing relatively few false positives.

| **parameters** | **values used in this study** |
|---|---|
| ibd_in | 1e-5 |
| ibd_out | 1e-5 |
| posterior probability cutoff for IBD1 | 0.99 |
| posterior probability cutoff for IBD2 | 0.8 |
| minimum segment length for IBD1 | 4 cM |
| minimum segment length for IBD2 | 2 cM |

**Table S3**: Percentage of IBD > 4 cM in grandparent-offspring and half siblings simulated with a human sex-specific genetic map. We used *ped-sim*[3] with a human sex-specific genetic map[4] and crossover interference model[5] to simulate 1,000 dyads of paternal and maternal grandparent-offspring and half sibling dyads each.

| | paternal | maternal |
|---|---|---|
| grandparent-offspring | 97.4% | 99.2% |
| half siblings | 96.5% | 99.2% |

**Table S4**: List of all dyads included in the final dataset. *Kinlabel* describes the assigned primary kinclass (i.e. closest consanguine relationship). The pedigree-based information includes relatedness estimates ($r_{PED}$), pedigree kin classes ($kinclass_{PED}$, entries separated by /@/) and information if the kinship is due to two common ancestors (e.g. full siblings) or one common ancestor (e.g. half siblings; $full\text{-}half_{PED}$, entries separated by /@/; see [6] for more details of the pedigree-based estimates). The IBD-based information includes estimates of global IBD sharing ($r_{IBD}$) and the number (*n IBD*) and length (*sum IBD*) of four different size classes (≥ 4, 8, 12, 20 cM respectively).

**Table S5:** Components to calculate the emission probability for the IBD2 state. We implemented this model in *ancIBD*[2].

| **g** | p(**g**) | p(**g**\|D) | p(**g**\|s=6) |
|---|---|---|---|
| (0,0,0,0) | $(1-p)^4$ | $(1-x_{1A})(1-x_{1B})(1-x_{2A})(1-x_{2B})$ | $(1-p)^2$ |
| (1,0,1,0) | $p^2(1-p)^2$ | $x_{1A}(1-x_{1B})x_{2A}(1-x_{2B})$ | $p(1-p)$ |
| (0,1,0,1) | $p^2(1-p)^2$ | $(1-x_{1A})x_{1B}(1-x_{2A})x_{2B}$ | $p(1-p)$ |
| (1,1,1,1) | $p^4$ | $x_{1A}x_{1B}x_{2A}x_{2B}$ | $p^2$ |

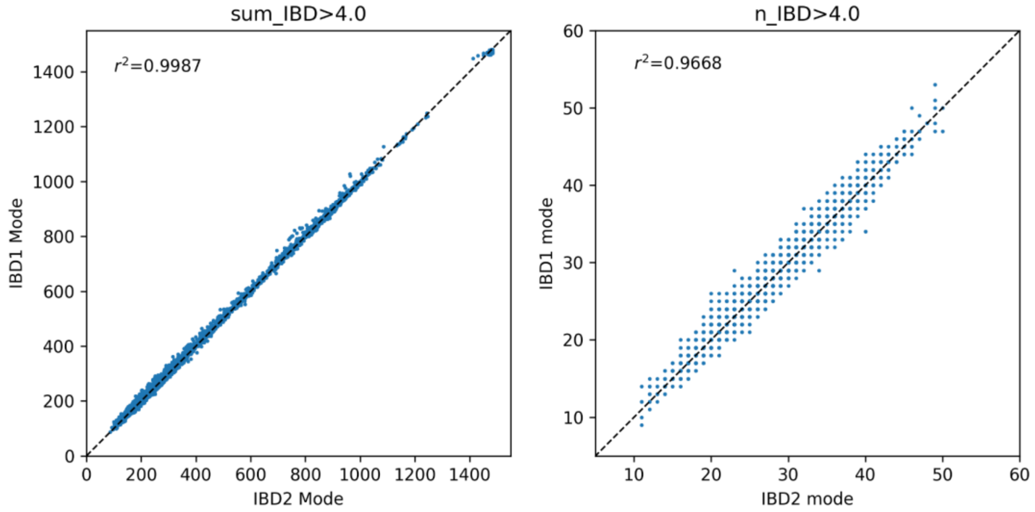


**Figure S1: IBD2 detection does not interfere with IBD1 detection.** Setting *ancIBD* to detect IBD2 segments as well as IBD1 segments does not substantively alter the detection of the overall amount of the genome IBD between dyads (left: sumIBD > 4 cM) or the number of IBD segments detected (right: nIBD > 4 cM). Estimates based on running *ancIBD* for IBD1 and IBD2 detection are shown on the x-axis; estimates based on running *ancIBD* for IBD1 detection only are shown on the y-axis.

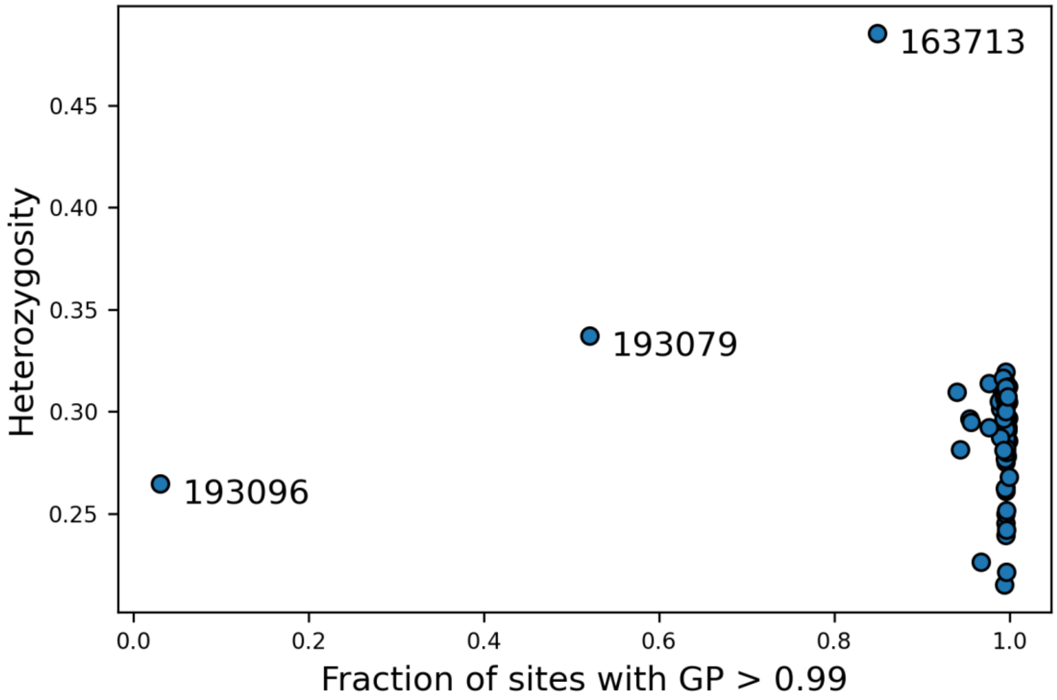


**Figure S2: Quality check of imputed samples.** Heterozygosity and fraction of sites with maximum GP > 99% for imputed samples. We define maxGP99fraction as the fraction of imputed sites for which the maximum genotype probability exceeds 99%. Most samples have maxGP99fraction > 95% and have heterozygosity levels around 28%. Three samples are outliers from the main cluster (due to very low coverage and/or sample contamination) and were excluded from any downstream analysis.

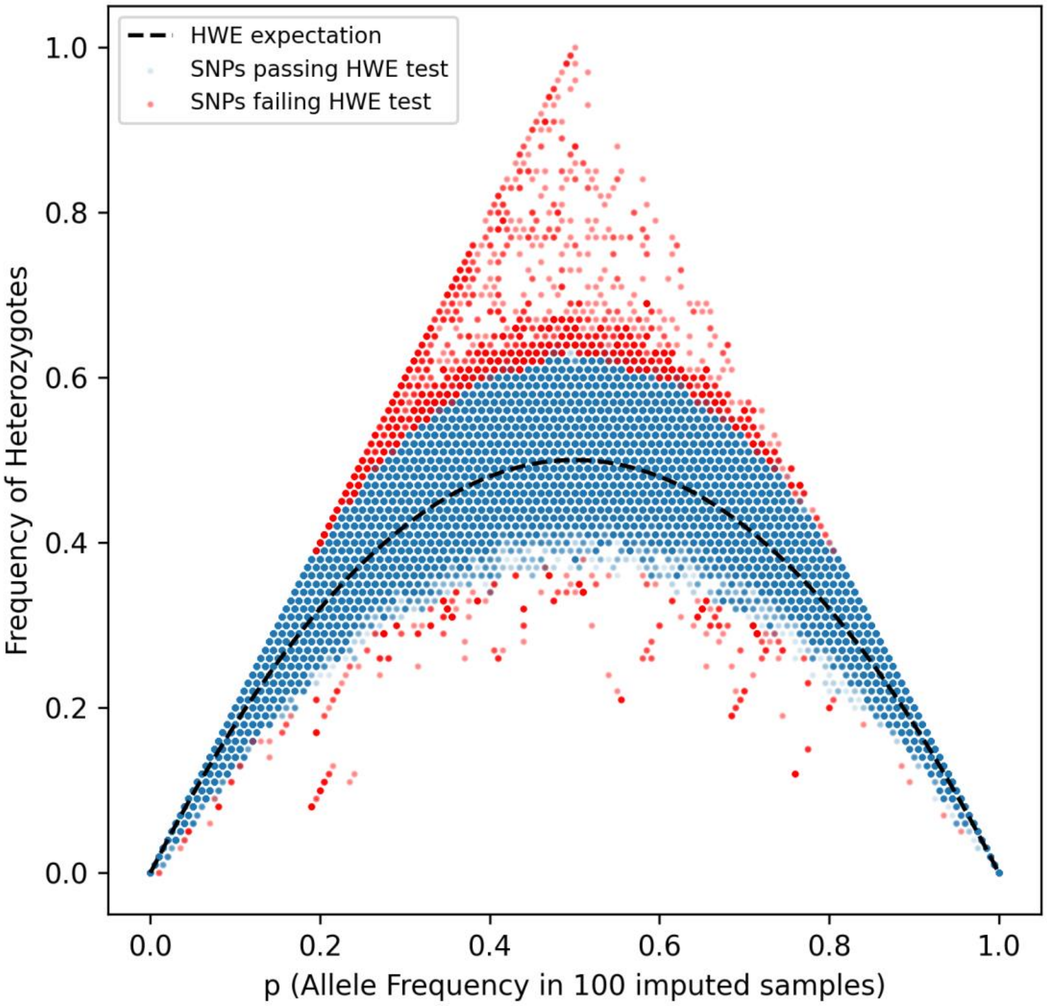


**Figure S3: Allele frequency and frequency of heterozygotes in the 100 imputed samples.** Under Hardy-Weinberg equilibrium (HWE), heterozygotes should occur at frequency $2p(1-p)$ for allele frequency $p$ (indicated by black dashed line). Blue dots are SNPs passing the Hardy-Weinberg equilibrium test implemented in *plink2* at a significance level of 0.01, and red dots are SNPs failing this test, which were subsequently excluded from IBD calling. This figure includes 2 individuals who we excluded from subsequent analysis due to uncertainty about their true identity; however, this type of sample labelling error should not affect population estimates of HWE.

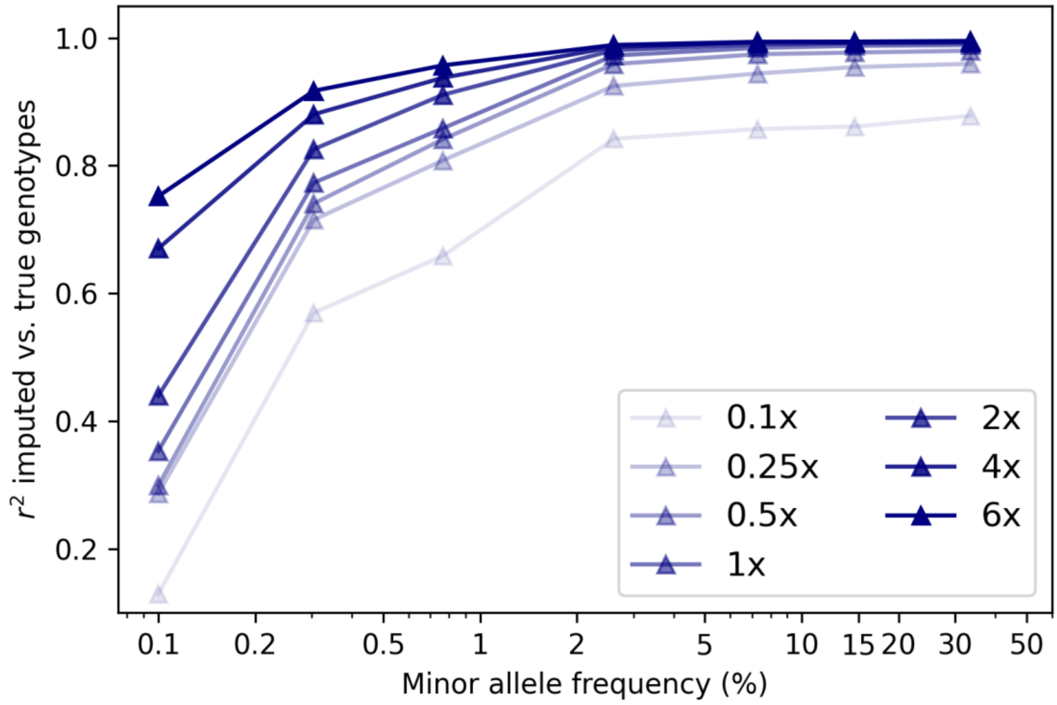


**Figure S4**: **Example imputation accuracy across various allele frequency bins and sequencing depths.** Imputation accuracy was measured by Pearson $r^2$ (y-axis), calculated in genotypes called from high-coverage data from individual 134,814 and genotypes called from downsampled data. We used the *GLIMPSE_concordance* utility (a submodule of *GLIMPSE*) for computing Pearson $r^2$ in allele frequency bins of [0.001, 0.002, 0.005, 0.01, 0.05, 0.1, 0.2, 0.5]. Each triangle's x-axis is the average allele frequency of all variants falling into that allele frequency mean.

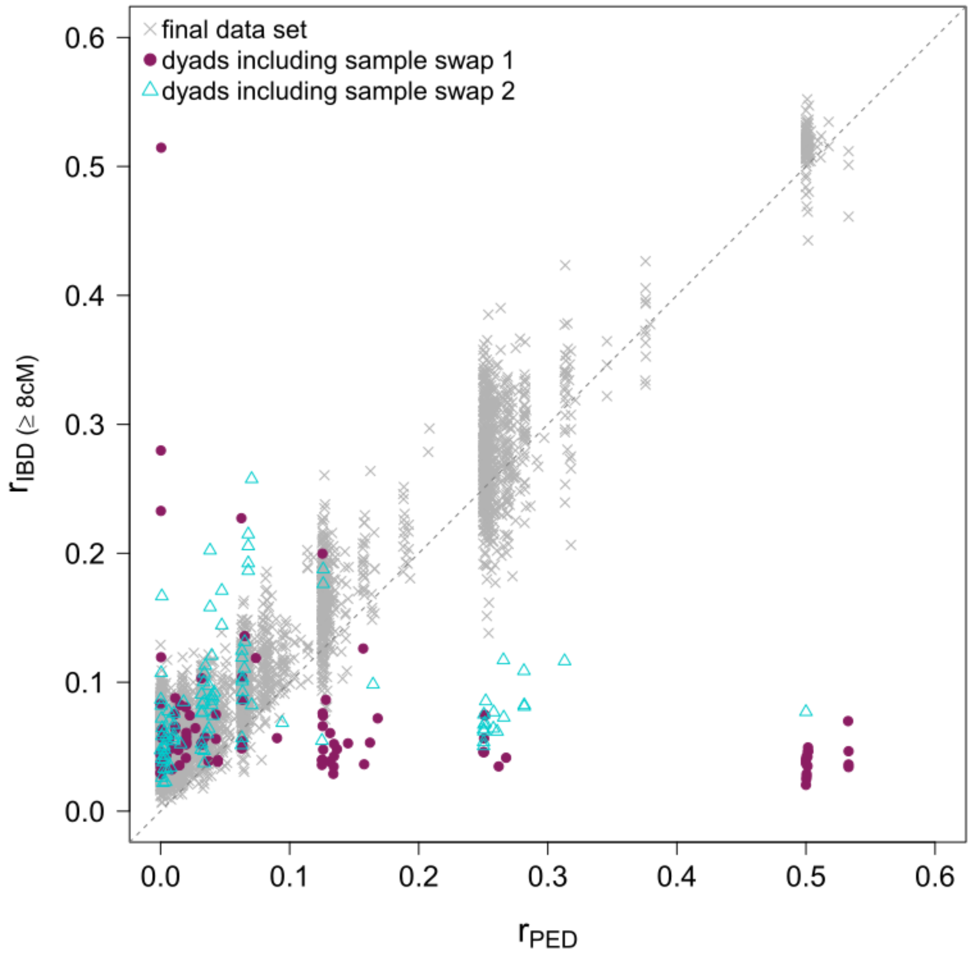


**Figure S5: Identification of samples of uncertain identity.** Comparison between IBD- and pedigree-based relatedness estimates. Gray crosses show the final dataset we used in this study. Colored symbols show dyads including two individuals who were removed from most analyses because their true IDs are uncertain. The gray dashed line demonstrates the perfect correlation between the two variables.

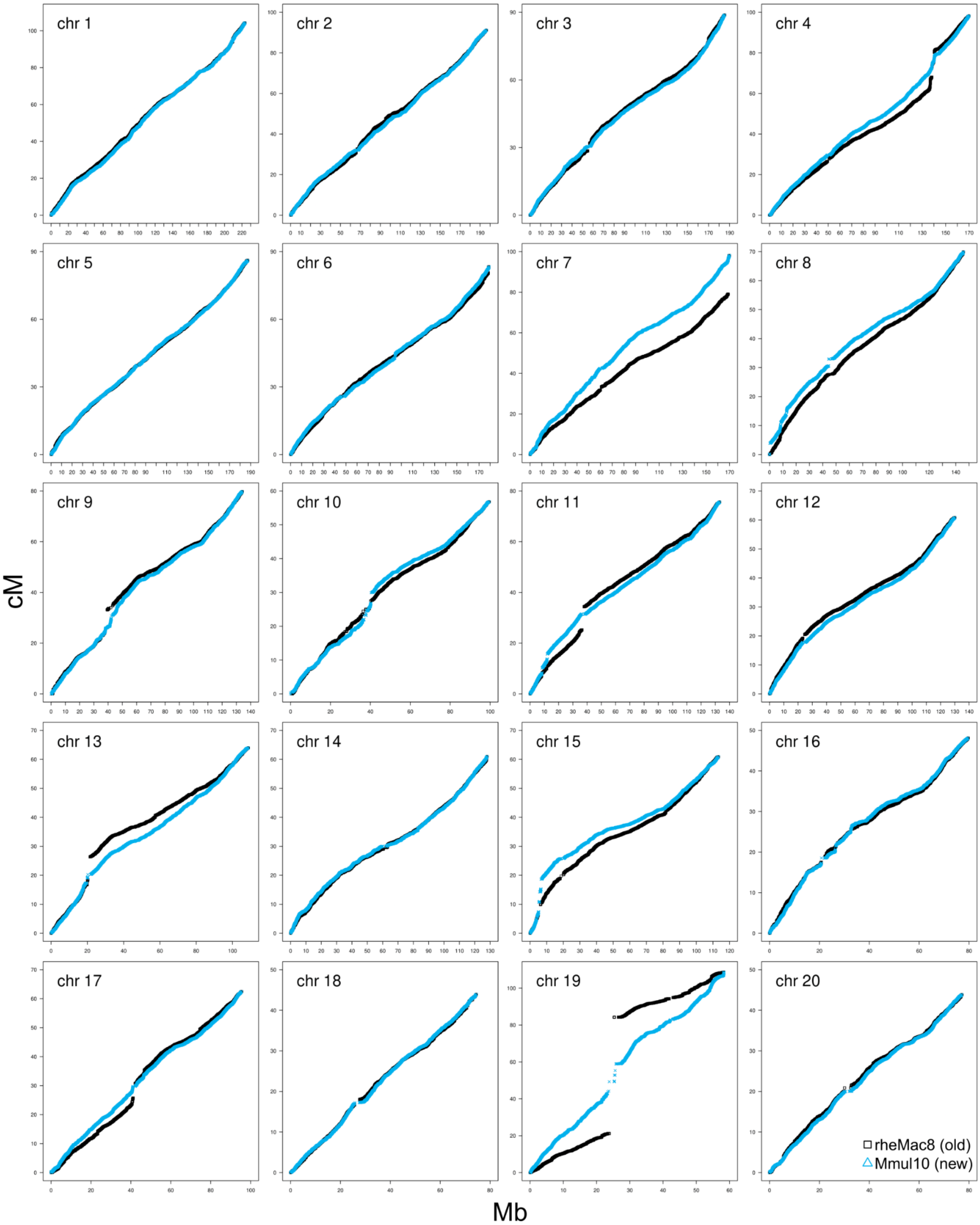


**Figure S6: Comparison between recombination maps.** The previously established recombination map is based on the *rheMac8* assembly, while the new one we presented here is based on the *Mmul10* assembly. To make a comparison between the two maps possible, we lifted over the coordinates from *rheMac8* to *Mmul10* before plotting them. In some chromosomes (e.g. 7 and 19) relatively large gaps are visible in the *rheMac8* map. Due to filtering out SNPs with unusually high linkage disequilibrium (see Methods for details), these gaps are far less pronounced in the *Mmul10* map.

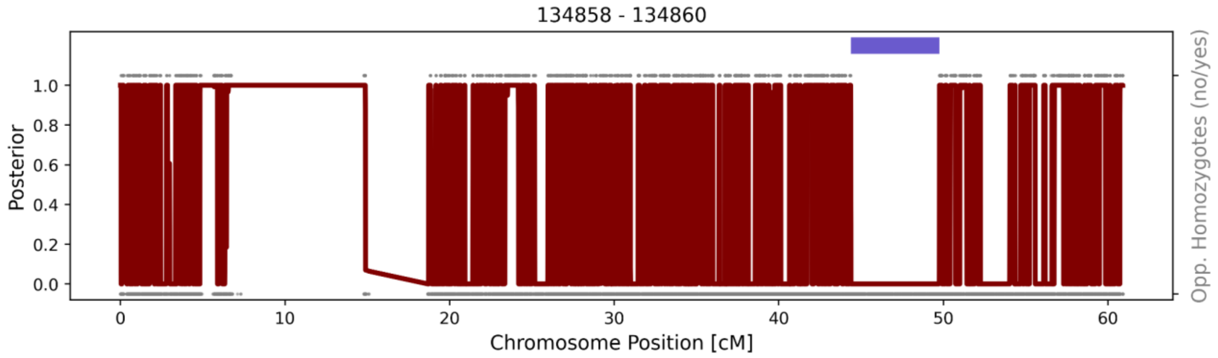


**Figure S7: Regions on chromosome 15 with few or no SNPs.** The regions with missing SNPs are ~7.31 cM to ~18.68 cM long. We masked this and other similar regions (a total of ~40cM) when calculating precision and recall for detected IBD segments. We visualize the *ancIBD* results on a dyad from animals 134858 and 134860 downsampled to 1× coverage. The gray dots at the very bottom indicate the locations of all SNPs. The red lines depict the posterior probability of not being in an IBD state. The gray dots at the top indicate the locations of opposing homozygotes. The horizontal blue bar indicates an inferred IBD segment.

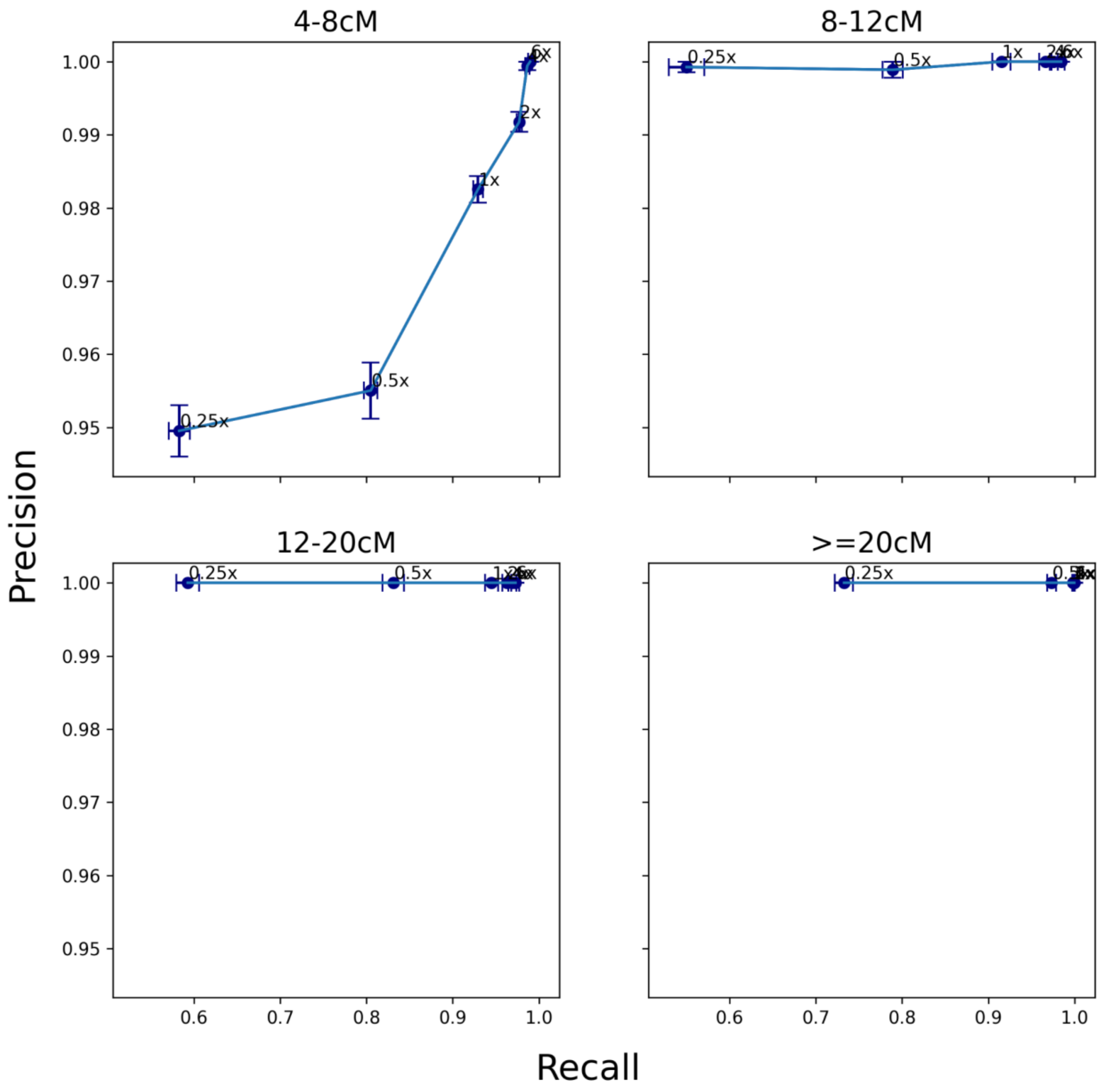


**Figure S8: Precision and recall of IBD1 after downsampling and imputation.** We computed precision and recall of IBD1 segments in four length bins. Average values are computed from 25 independent downsampling replicates. The error bars represent one standard error.

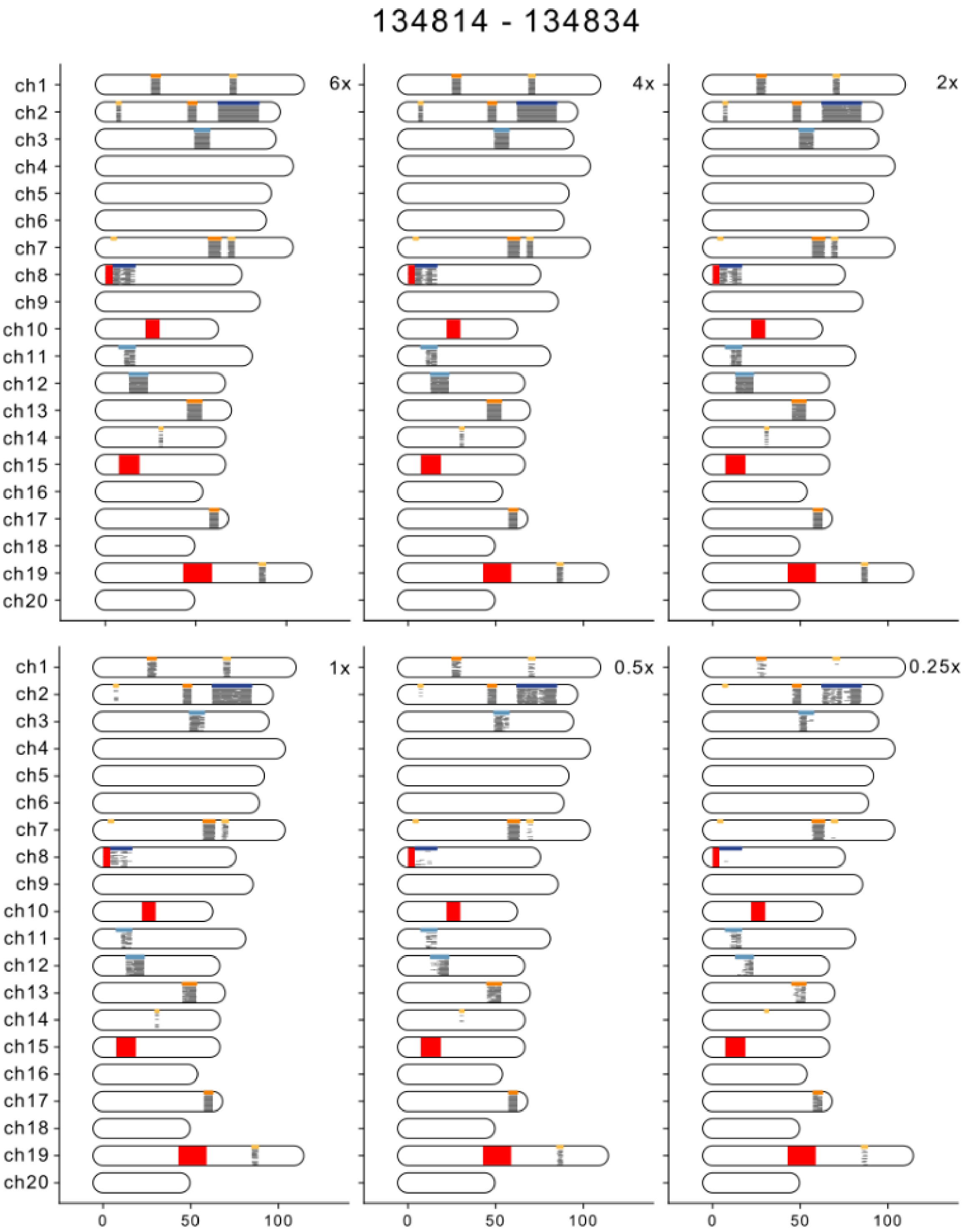
134814 - 134834
6x
4x
2x
1x
0.5x
0.25x
ch1
ch2
ch3
ch4
ch5
ch6
ch7
ch8
ch9
ch10
ch11
ch12
ch13
ch14
ch15
ch16
ch17
ch18
ch19
ch20
0
50
100

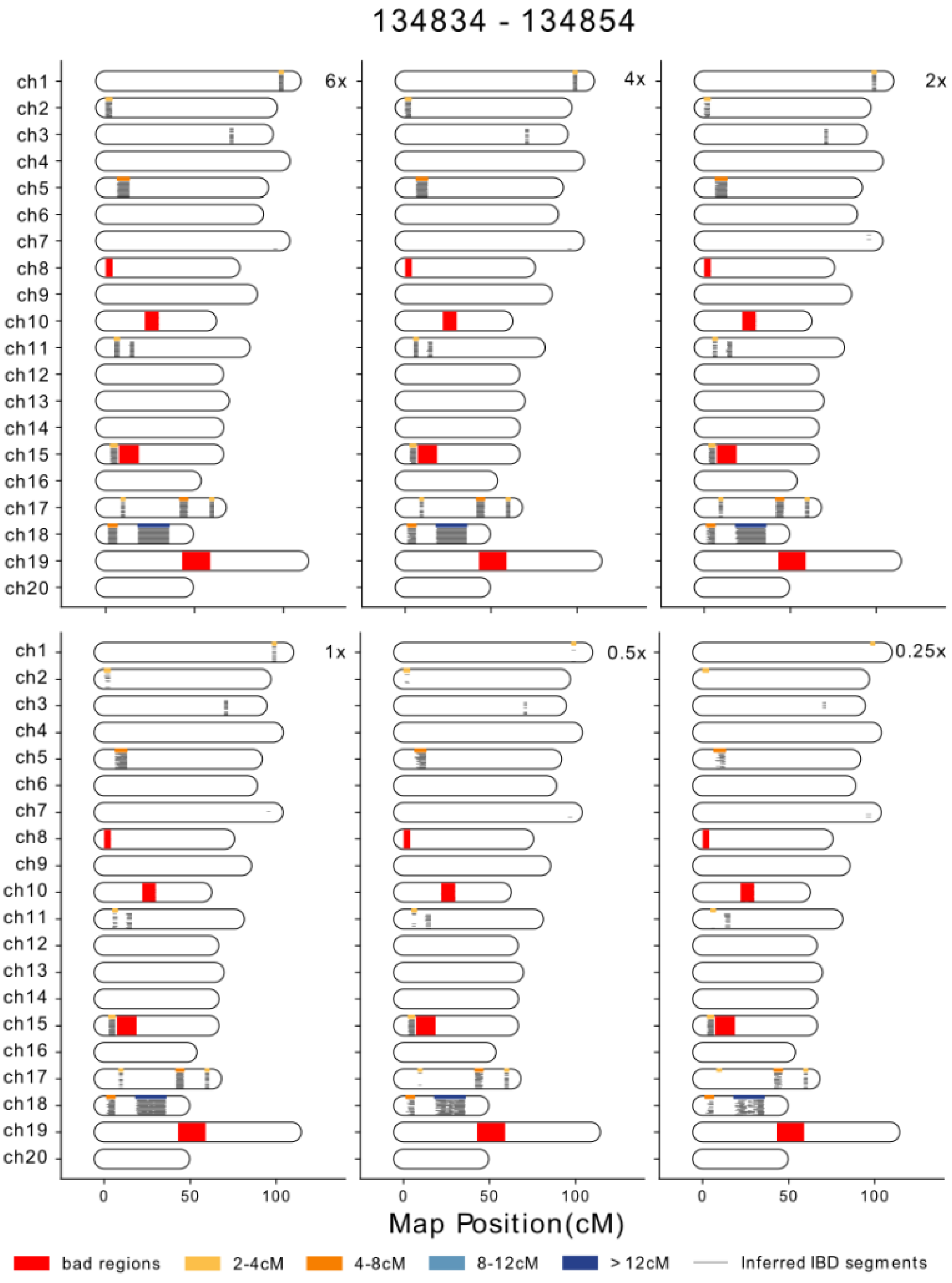


**Figure S9: Inferred and ground truth IBD2 segments.** Data are shown for the dyads 134814 – 134834 (top) and 134834 – 134854 (bottom). Ground truth IBD2 (called by IBIS on the high-coverage data) are depicted as thick bars at the top of each chromosome, and colored differently according to their length. IBD2 inferred from downsampled data (from 6× to 0.25×, with higher coverage data sets to the left) are visualized as thin black lines stacked together. Each replicate occupies one horizontal space. Regions with few SNPs are masked with thick brown bars.

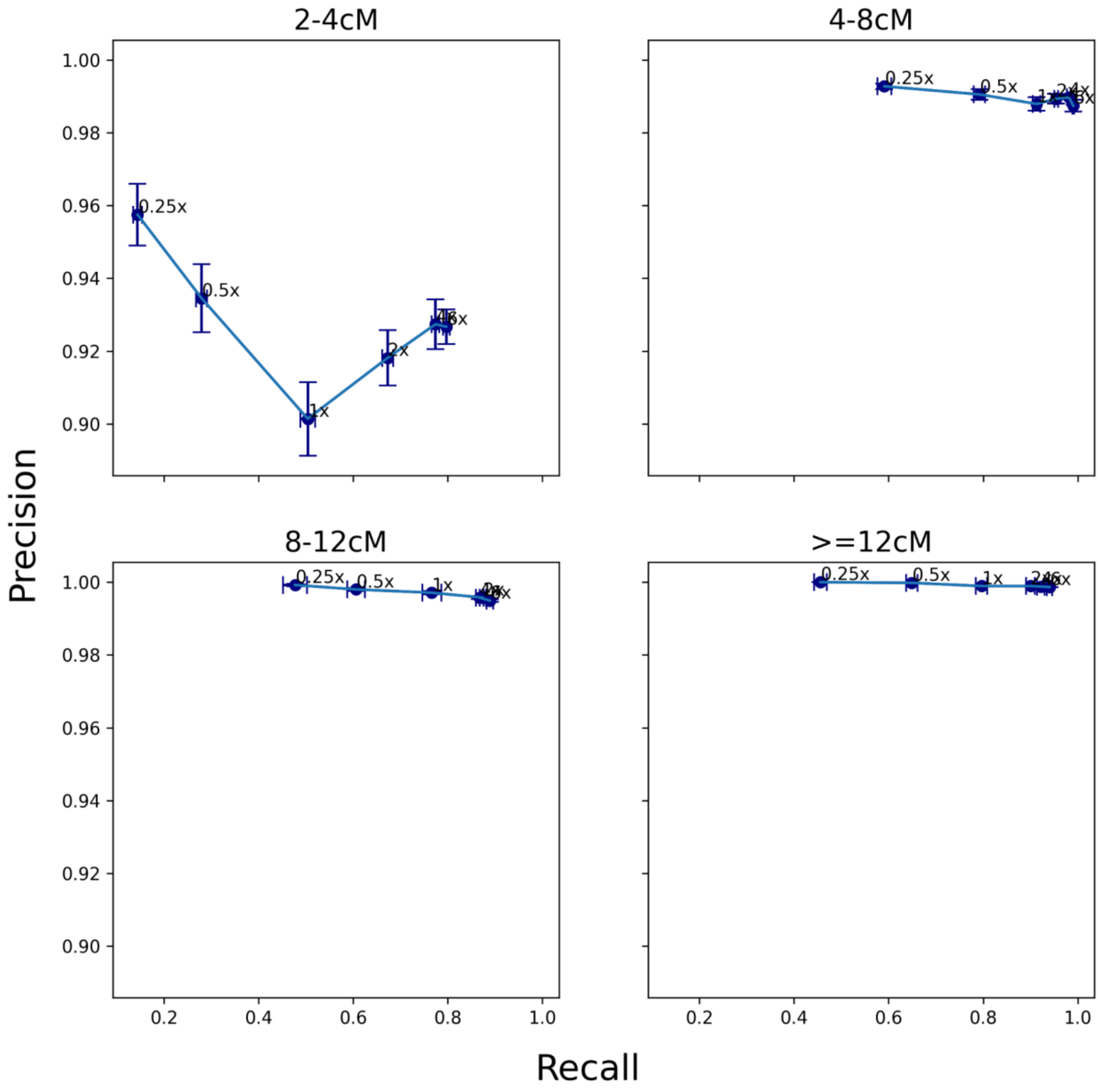


**Figure S10: Precision and recall of IBD2 after downsampling and imputation.** We computed precision and recall of IBD2 segments in four length bins. Average values are computed from 25 independent downsampling replicates. The error bars represent one standard error.

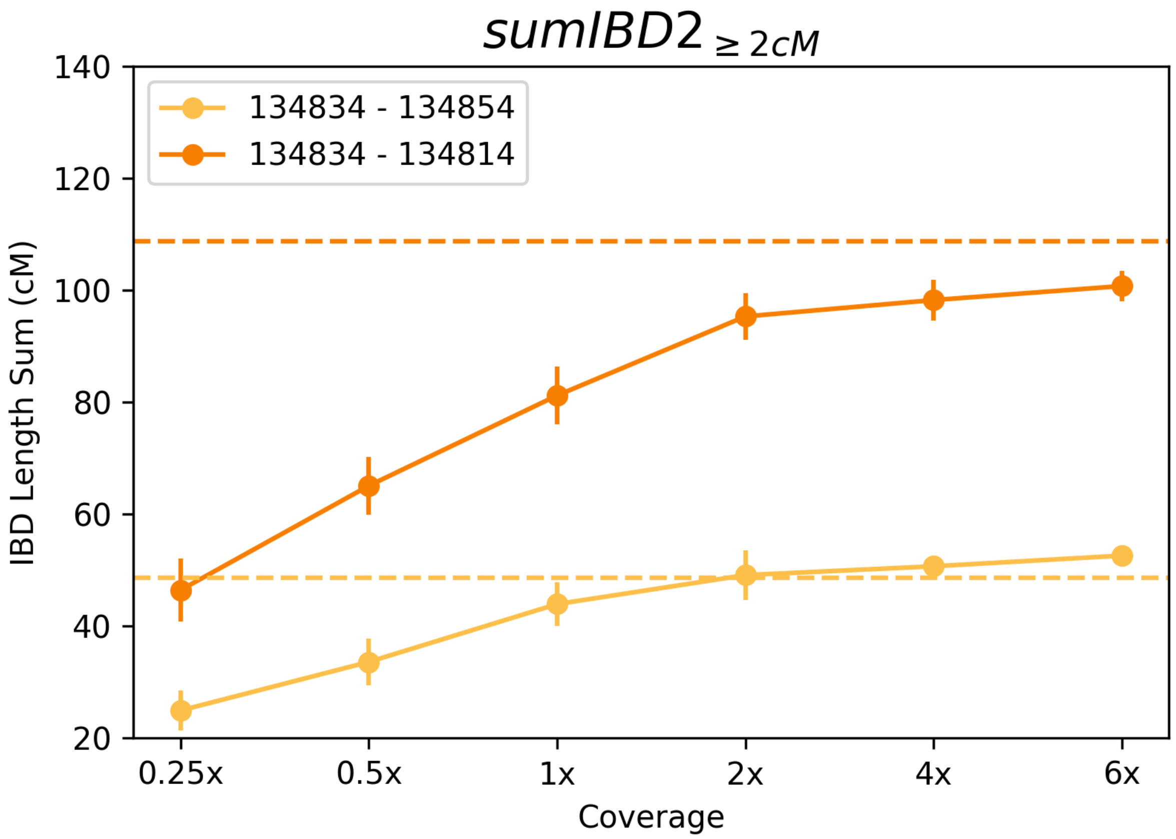


**Figure S11: Dependency of total inferred IBD2 length on coverage.** $sumIBD2_{\geq 2cM}$ as a function of coverage for two dyads with high-coverage data. The error bar indicates one standard deviation, calculated from 25 independent downsampling of the original high-coverage data. The dashed horizontal lines indicate ground truth $sumIBD2_{\geq 2cM}$, obtained from calling IBD2 in the high-coverage data using *IBIS*.

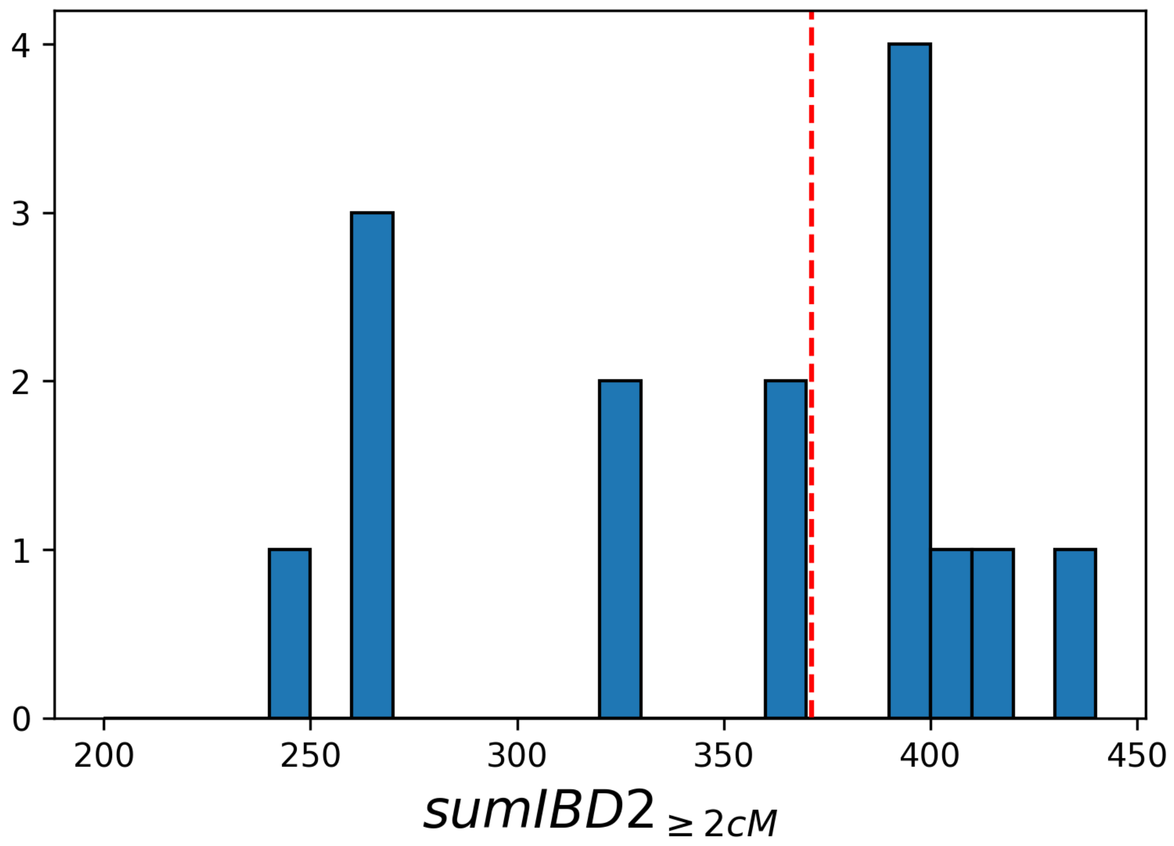


**Figure S12: Histogram of sumIBD2$_{\geq 2cM}$ for full siblings.** Data are shown for all full siblings in the imputed data set. The red vertical line indicates the expected sumIBD2 for full sibling dyads, which is one-fourth of the total genome length.

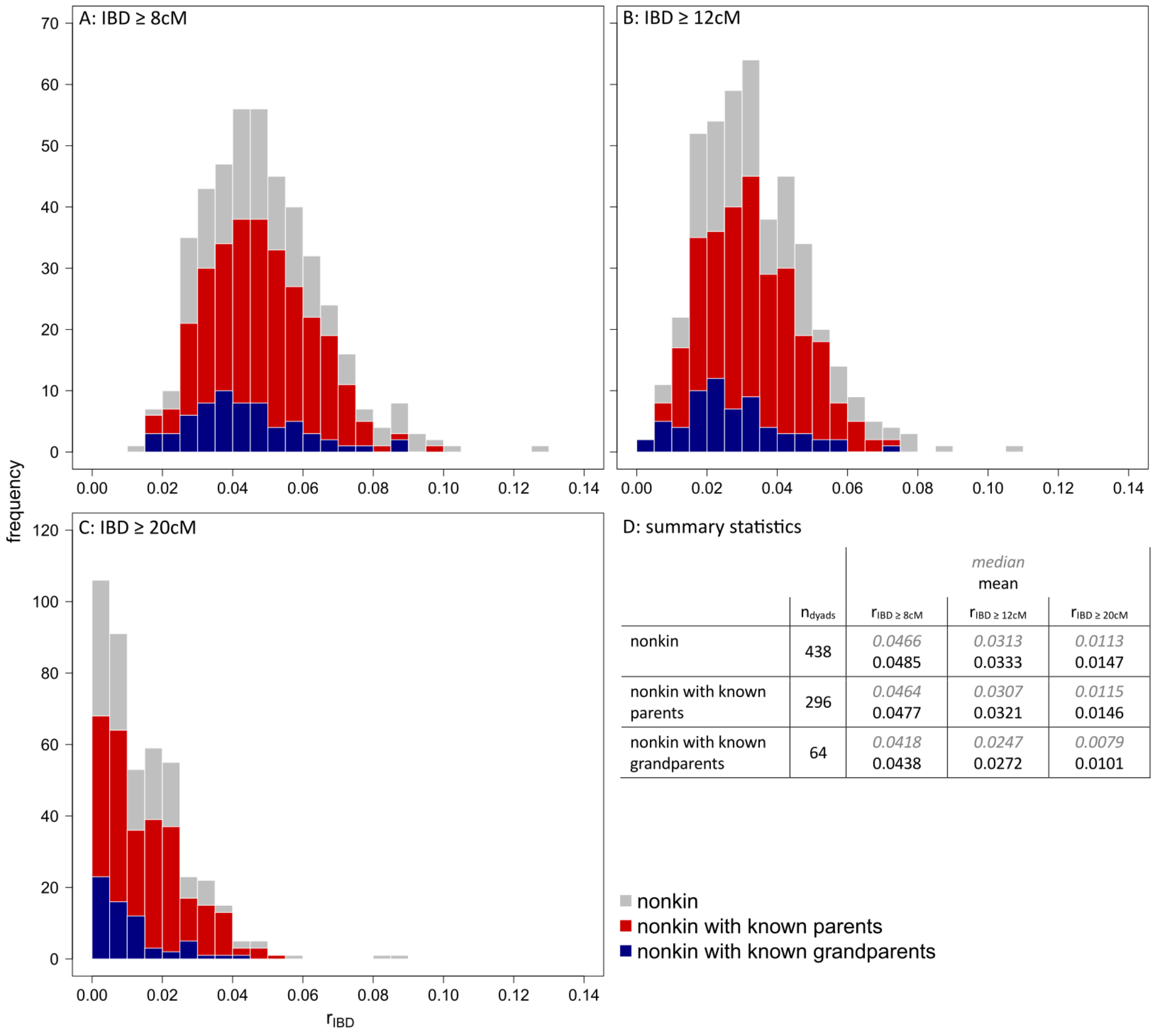


| | $n_{dyads}$ | $r_{IBD \geq 8cM}$ | $r_{IBD \geq 12cM}$ | $r_{IBD \geq 20cM}$ |
|---|---|---|---|---|
| | | *median* / mean | | |
| nonkin | 438 | *0.0466* 0.0485 | *0.0313* 0.0333 | *0.0113* 0.0147 |
| nonkin with known parents | 296 | *0.0464* 0.0477 | *0.0307* 0.0321 | *0.0115* 0.0146 |
| nonkin with known grandparents | 64 | *0.0418* 0.0438 | *0.0247* 0.0272 | *0.0079* 0.0101 |

**Figure S13: Histogram of background relatedness.** Gray bars show all dyads that are classified as nonkin by the pedigree (gray). Colored bars show subsets excluding dyads for which at least one individual's parent (red) and/or grandparent (blue) is unknown. (A) shows IBD segments ≥ 8cM, (B) ≥ 12cM and (C) ≥ 20cM. (D) shows sample sizes, medians and means of each subset.
